# *LY6S,* a New Interferon-Inducible Human Member of the Ly6a-Subfamily Expressed by Spleen Cells and Associated with Inflammation and Viral Resistance

**DOI:** 10.1101/2021.12.16.472998

**Authors:** Moriya Shmerling, Michael Chalik, Nechama I. Smorodinsky, Alan Meeker, Sujayita Roy, Orit Sagi-Assif, Tsipi Meshel, Artem Danilevsky, Noam Shomron, Shmuel Levinger, Bar Nishry, David Baruchi, Avital Shargorodsky, Ravit Ziv, Avital Sarusi-Portuguez, Maoz Lahav, Marcelo Ehrlich, Bryony Braschi, Elspeth Bruford, Isaac P. Witz, Daniel H. Wreschner

**Author notes:** Corresponding authors. Daniel H. Wreschner Isaac P. Witz.

## Abstract

Syntenic genomic loci on human chromosome 8 (hChr8) and mouse chromosome 15 (mChr15) code for LY6/Ly6 (lymphocyte antigen 6) family proteins. The 23 murine *Ly6* family genes include eight genes that are flanked by the murine *Ly6e* and *Ly6l* genes and form an Ly6 subgroup referred to here as the Ly6a subfamily gene cluster. *Ly6a,* also known as *Sca1* (*Stem Cell Antigen-1*) and *TAP* (*T-cell activating protein*), is a member of the Ly6a subfamily gene cluster. No *LY6* genes have been annotated within the syntenic *LY6E* to *LY6L* human locus. We report here on *LY6S*, a solitary human *LY6* gene that is syntenic with the murine Ly6a subfamily gene cluster, and with which it shares a common ancestry. *LY6S* codes for the interferon-inducible GPI-linked LY6S-iso1 protein that contains only 9 of the 10 consensus LY6 cysteine residues and is most highly expressed in a non-classical cell population. Its expression leads to distinct shifts in patterns of gene expression, particularly of genes coding for inflammatory and immune response proteins, and LY6S-iso1 expressing cells show increased resistance to viral infection. Our findings reveal the presence of a previously un-annotated human interferon-stimulated gene, *LY6S,* which has a one to eight ortholog relationship with the genes of the Ly6a subfamily gene cluster, is most highly expressed in spleen cells of a non-classical cell-lineage and whose expression induces viral resistance and is associated with an inflammatory phenotype and with the activation of genes that regulate immune responses.

**One Sentence Summary:** *LY6S* is a newly discovered human interferon-inducible gene associated with inflammation and with resistance to viral replication.

## Introduction

The human genome contains at least 48 genes and the mouse genome at least 61 genes, all coding for proteins that contain a signature pattern of 10 cysteine residues, resulting in a distinctive disulfide bonding pattern. Originally discovered as GPI-linked cell-surface proteins present on lymphocytes (LY (*1*)), they are now known to be expressed by diverse cell types(*2-4*). They typically form a three-fingered protein (TFP) and contain a domain found in the urokinase-type plasminogen activator receptor protein (uPAR, encoded by *PLAUR*) and are also referred to as LY6/uPAR or LU (Lymphocyte antigen 6-uPAR) proteins. These proteins locate extracellularly either linked to the cell membrane via a glycosyl-phosphatidyl-inositol (GPI) anchor, as transmembrane proteins or as fully secreted proteins as seen for example in the PATE proteins (*5*). The transmembrane proteins include the receptors for proteins belonging to the transforming growth factor-beta family, such as BMP2 and activin (*6*). CD59 encodes both a GPI-anchored and a secreted LY6 family protein. Only a few of the TFP/LY6/uPAR proteins have known functions, with the most well characterized being CD59 (*7, 8*), uPAR/PLAUR (*9*) and GPIHBP1 (*10*). A common theme for these proteins is that the LY6 domain mediates interactions with other proteins. This is exemplified by CD59, a protein that regulates complement-mediated cell lysis by binding to the complement membrane attack complex, via uPAR, where 3 Ly6 domains form the binding complex for the serine protease urokinase-type plasminogen (*11*), and by GPIHBP1, an endothelial cell protein that binds lipoprotein lipase and transports it to the capillary lumen (*10, 12*). The variety of functions both known and postulated for the LY6 family proteins suggests that nature has usurped the skeleton scaffold of the TFP fold in order to execute a large number of diverse activities. Indeed, among the various LY6 proteins there are large variations in hydrophobicity levels and isoelectric points, in addition to low overall sequence similarity (*2-4, 13*).

Mouse chromosome 15 harbors a locus replete with 23 *Ly6* family genes (Fig. 1), most of which code for Ly6 proteins linked to the cell surface by a GPI anchor. By utilizing alternative splicing mechanisms some of these genes also code for secreted Ly6 proteins.

**Fig. 1.**
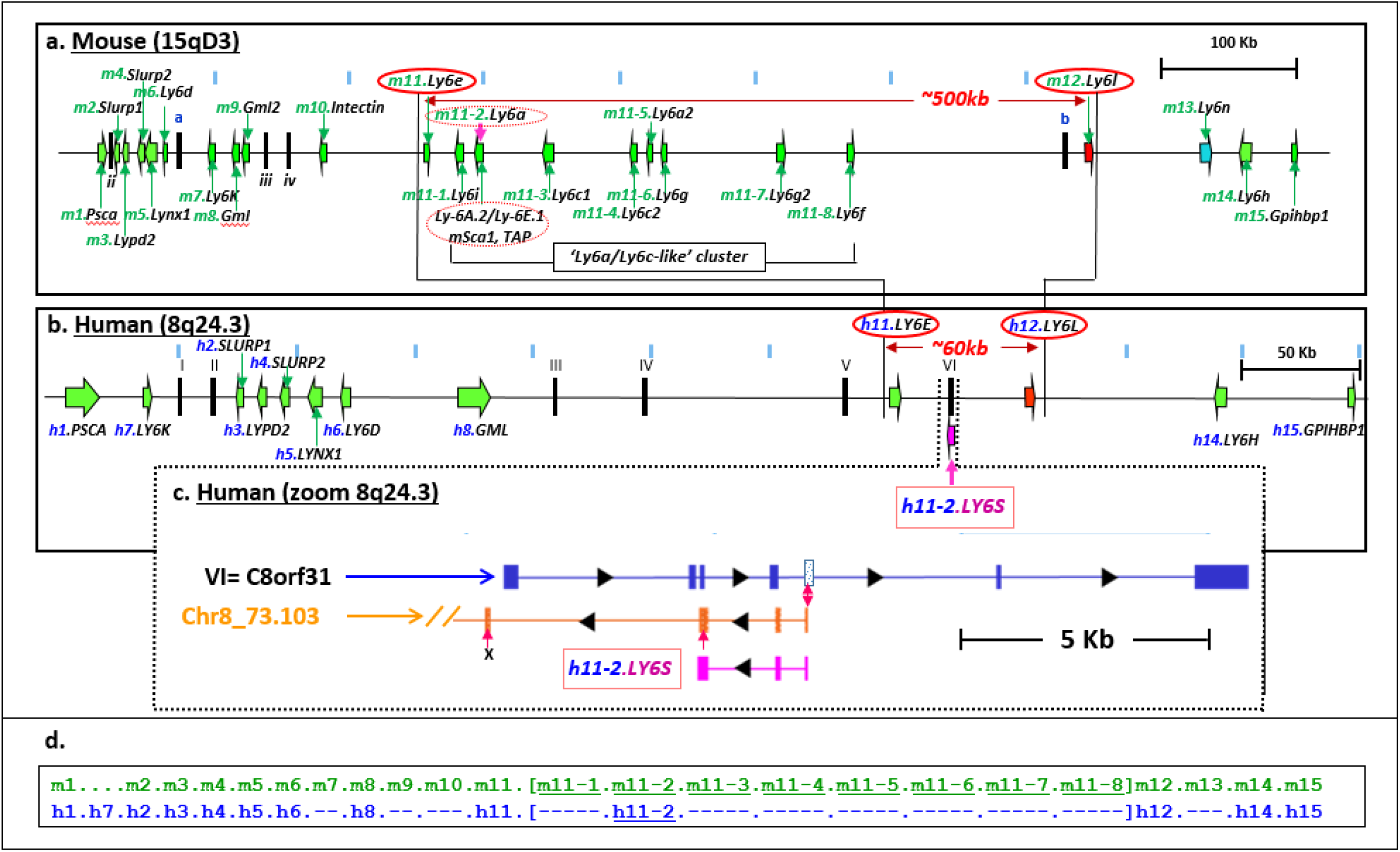
Ly6/LY6 loci in mouse chr15qD3 and human chr8q24.3 and location of human LY6S. All known *Ly6/LY6* genes located on murine chr15qD3 and human chr8q24.3 are shown as green arrows with transcriptional orientation as indicated (panels a and b, respectively). Non-Ly6/LY6 genes appear as vertical black bars, and are marked by small and capital Roman numerals (mouse and human, respectively). The murine *Ly6* genes in addition to their accepted designations are indicated as m1 through to m15 and the 8 murine genes belonging to the Ly6a subfamily cluster are indicated by m11-1 through m11-8; the corresponding human genes have the prefix ‘h’, and *LY6S* is designated h11-2 (panels a, b and d). The murine *Ly6e* and *Ly6l* genes flanking the Ly6a subfamily cluster and their corresponding human *LY6E* and *LY6L* orthologs are encircled by red ovals (panels a and b). A zoom-in of the region in which human *C8orf31* and the Genscan predicted gene (in orange) reside is shown in panel c. Panel d-Schematic comparison of the murine (m1-m15) and corresponding human (h) genes-dashes indicate no known human homolog.

One of the most widely investigated *Ly6* genes is *Ly6a,* which is located within the chromosome 15 locus (*14*) and codes for an interferon-inducible GPI-linked protein, also known as TAP (T-cell Activating Protein) and stem cell antigen-1 (Sca1). *Ly6a* is upregulated by interferons and is implicated in the activation of T cells, as well as in inflammatory and immune-related processes. Similarly, the murine genes *Ly6c* and *Ly6g* which are also in this cluster are used extensively to analyze cells of the myeloid lineage, especially of the mouse spleen (for example (*15-17*)) and *Ly6c* expression is used as a marker for murine blood monocytes (*18, 19*). Despite the hundreds of publications on each of these three murine genes, many focusing on their role in inflammation, neither a putative human ortholog(s) nor their actual functions have to date been reported.

Investigation of the mouse chromosome 15 *Ly6* locus and its comparison with the human chromosome 8 syntenic counterpart (Fig. 1) revealed that *Ly6a* locates to a 500kb sub-region within the mChr15 Ly6 locus, flanked upstream and downstream by the *Ly6e* and *Ly6l* genes, and contains a total of 8 Ly6 family genes. Many of these genes, if not all, are inducible by interferons and code for proteins that are markedly upregulated in inflammatory processes and immune-related diseases (*20-22*).

The phylogenetic tree analysis of the human Chr 8 and murine Chr5 *LY6* loci (Fig. 1) shows that the 8 Ly6 family genes located within the 500kb sub-region, group together in a clade with high bootstrap support and are more distantly related to the *Ly6* genes located outside of the *Ly6e* to *Ly6l* locus. Cumulatively, this indicates that the 8 genes within the *Ly6e* to *Ly6l* locus form a Ly6 family subgroup, referred to here as the Ly6a subfamily. It is surprising, therefore, that no human homolog(s) to this subfamily has been previously identified.

In contrast to the ∼500kb murine *Ly6a* subfamily locus, the corresponding syntenic human region also flanked by *LY6E* and *LY6L* is only ∼ 60kb (Fig. 1) and contains no annotated human LY6 genes. We report here the identification of a new human LY6 family gene designated *LY6S* by the HUGO Gene Nomenclature Committee (HGNC) (*23*) within this region. *LY6S* codes for a LY6 protein that phylogenetic analyses (Fig. 2) indicate has a 1 to 8 orthologous relationship with the murine *Ly6a* subfamily cluster. It codes for a membrane-linked LY6S protein that is most highly expressed in the human spleen by a non-classical human spleen cell and is associated with the regulation of cell growth, inflammatory and immune-related responses, and with resistance to infection by certain viruses.

**Fig. 2.**
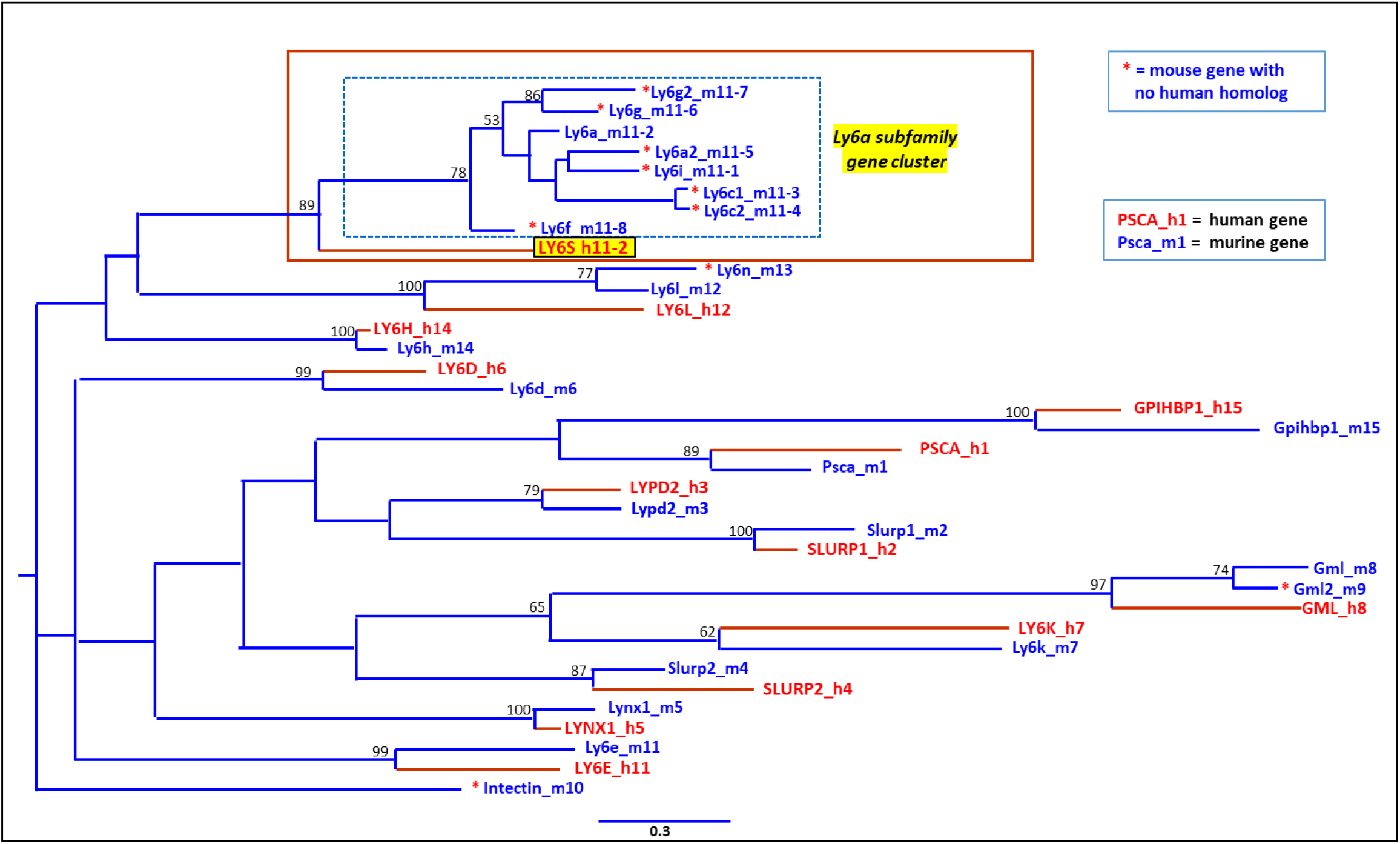
Phylogenetic tree of the known murine/human Ly6/LY6 genes located on mouse chr15qD3 and human chr8q24.3. Maximum likelihood tree constructed as detailed in Materials and Methods. Murine and human *Ly6/LY6* genes are indicated by blue and red fonts respectively. Murine genes belonging to the Ly6a subfamily are bordered by the dashed blue-lined rectangle, and the human *LY6S* gene is shown in red fonts against a yellow background. Bootstrap support values >50% are shown above the nodes, branch lengths represent the expected number of substitutions per site and the tree is shown as unrooted. The red asterisks indicate mouse genes with no previously identified human ortholog.

## Results

### Analysis of mChr15 and hChr8 Ly6/LY6 loci reveals a new human LY6 gene syntenic with Ly6a

*Ly6a* resides on mouse chromosome 15 in a 900 kb genomic locus replete with other genes coding for Ly6 family proteins (Fig. 1). The syntenic human locus on chromosome 8 is also rich in LY6 genes, yet at about 540kb is considerably smaller than the corresponding mouse locus and contains far fewer LY6 family genes. The murine sub-region containing *Ly6a* is demarcated by the *Ly6e* and *Ly6l* genes, spans ∼500kb and contains 8 Ly6 genes. In contrast, the human *LY6E* and *LY6L* genes (Fig. 1, panel b) span a region of ∼60 kb within which no annotated LY6 genes reside. Because (a) the mouse *Ly6a* gene lies downstream to *Ly6e*, within the genomic neighborhood demarcated by the *Ly6e* and *Ly6l* genes and as (b) *LY6L* is predicted to be the human ortholog of *Ly6l* by multiple orthology resources (see the HGNC HCOP tool: https://www.genenames.org/data/gene-symbol-report/#!/hgnc_id/HGNC:52284), we expected that if a human ortholog of *Ly6a* were to exist, then it too should reside within the human syntenic segment bordered by *LY6E* and *LY6L* which spans only ∼60 kb (see Fig 1B). No annotated human LY6 genes appear in this region, yet GenScan predicts a gene (chr8_73.103, Fig. 1C) coding for an LY6 family protein that shows a high level of sequence homology to the mouse Ly6a protein. Furthermore, the *TransMap* cross-species alignment algorithm (*24, 25*) that maps genes and related annotations in one species versus another by using synteny-filtered pairwise genome alignments to determine the most likely ortholog, maps *Ly6a* at the same position as the GenScan predicted human LY6 gene (Fig. S1).

### Experimental confirmation for LY6S mRNAs

Neither ESTs nor any other experimental evidence support the GenScan chr8_73.103 gene. Notably, transcription of the LY6 gene predicted by GenScan (Fig. 1c) is in the opposite direction to that of a known gene, *C8orf31*. *C8orf31,* taxonomically restricted to primates, is transcribed from the complementary strand and shows extensive exon overlap with exons from the GenScan predicted human LY6 gene.

Therefore, forward and reverse primers for the predicted chr8_73.103 gene serve respectively for the *C8orf31* transcripts as reverse and forward primers, and our initial attempts to identify transcript(s) coded by the predicted chr8_73.103 gene yielded a mixture of RT-PCR products, apparently derived from both strands, that could not be directly sequenced. However, TOPO cloning of the PCR products revealed transcripts obviously derived from *C8orf31*. Several of these RT-PCR products contained an additional un-annotated *C8orf31* exon (flanked by consensus splice sites), that showed extensive overlap with the predicted first coding exon of chr8_73.103. This finding was particularly informative as it specified the location of all sequences, here defined as “poison” sequences, that completely overlap between *C8orf31* and the GenScan predicted Chr8_73.103 gene (Fig. 3 a, b and c).

**Fig 3.**
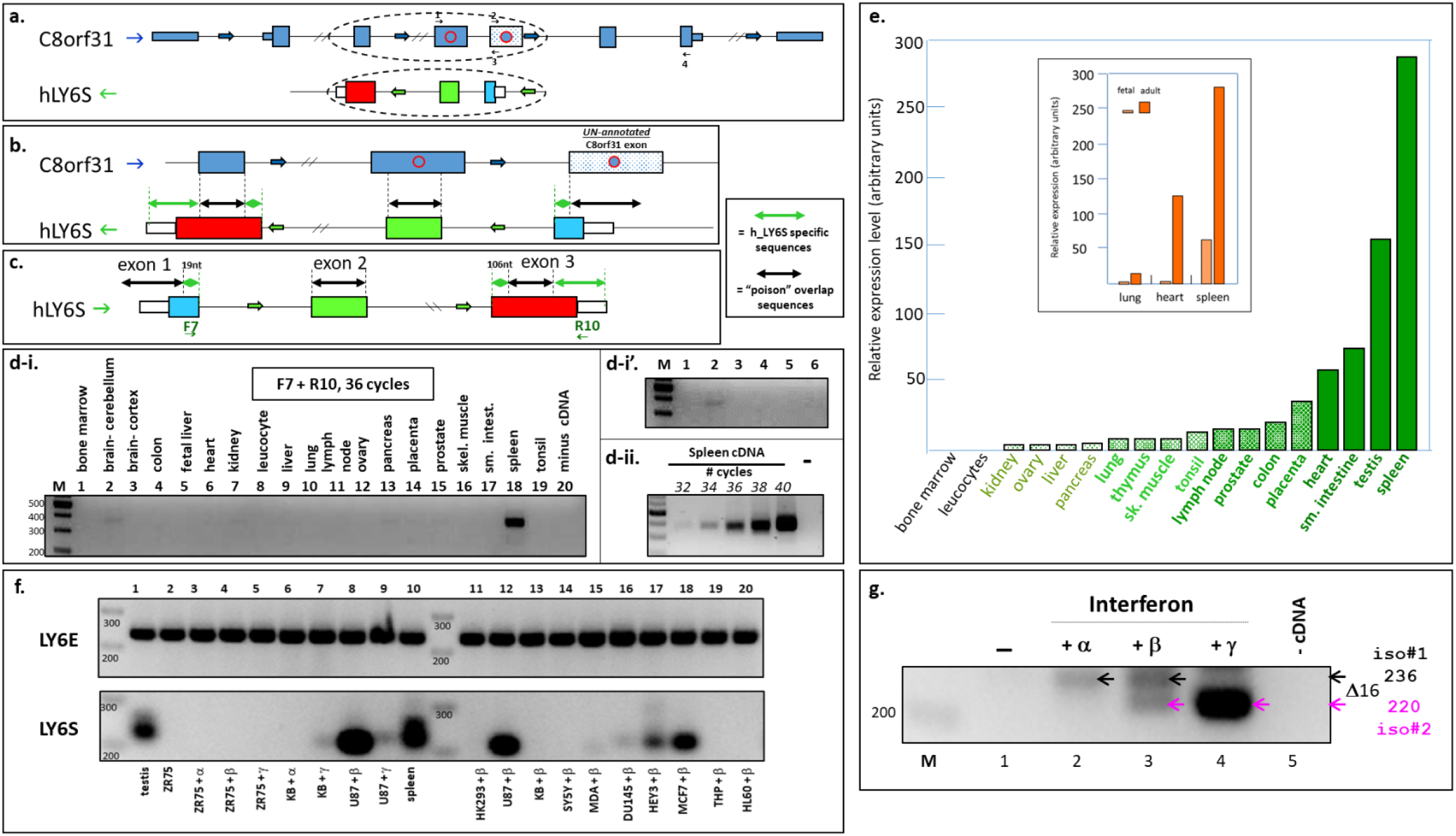
Exon overlap of the transcriptional units of LY6S and C8orf31, expression of LY6S mRNA in the human spleen and other tissues and induction of LY6S mRNA expression by interferons. **Panels a and b-***C8orf31* transcription proceeds left to right (blue arrows), whereas *LY6S* is transcribed from the opposite strand, right to left (green arrows). The un-annotated *C8orf31* exon is indicated by the stippled light blue box. Overlapping sequences are designated “poison” (black double-sided arrows), whereas regions specific for *LY6S* are marked by green double-sided arrows. Panel c-Transcription of *LY6S* is shown from left to right, together with the *LY6S* specific forward and reverse primers F7 and R10. Panels d-i, d-i’ and d-ii-RT-PCR analysis of cDNAs derived from human tissues (36 cycles). Higher contrast exposure of lanes 1-6 is shown in panel d-i’ and for different numbers of PCR cycles with spleen cDNA (panel d-ii). Panel e-*LY6S* expression in various adult tissues, as indicated, was assessed by qPCR. The inset compares relative LY6S expression levels in fetal and adult tissues. Panel f-*LY6E* and *LY6S* expression (upper and lower panels, respectively) in testis and spleen (lanes 1 and 10, respectively) and from various human cell lines treated with interferon, as indicated, were assessed by RT-PCR. **Panel g**-*LY6S* expression in melanoma (M12CB3) cells that were untreated or treated with interferon alpha, beta or gamma (lanes 1-4, respectively) was assessed by RT-PCR.

The primers (F7 and R10, see Fig. 3c and Table S1) are specific solely for the predicted Chr8_73.103 gene henceforth referred to as *LY6S,* a gene symbol approved by the HGNC. Use of these primers revealed in RT-PCR analyses a particularly prominent RT-PCR product in the spleen (Fig 3d-i and Fig. 3d-ii), which showed the highest expression (Fig. 3d-i and d-ii and Fig. 3e). In fetal tissues the spleen was also the highest *LY6S* expresser. Yet *LY6S* expression in all fetal tissues was considerably lower than that seen in the corresponding adult tissues (inset, Fig. 3e). In contrast to its high expression in the lymphoid-rich spleen, *LY6S* expression was not detected in peripheral blood leucocytes or in bone marrow [Fig. 3d-i, compare lanes 1 and 8 with lane 18 (bone marrow, leucocytes and spleen, respectively) and Fig. 3e]. In addition to spleen, *LY6S* expression was detected in brain tissue (Fig. 3d-i, lane 2 and at higher contrast in Fig. 3d-i’, lane 2). In line with these results, increasing the number of PCR cycles to 40 showed, as expected, highest expression in spleen followed by testis (Fig. S2a, lanes 14 and 16, respectively). Significant expression was also detected in the parietal lobe of the brain and in the spinal cord, and lower levels in the thalamus and temporal lobe (Fig. S2a, lanes 8, 13, 17 and 15 respectively). Because of the therapeutic implications of expression of LY6S in the human brain (see Discussion) and in order to confirm these results, this experiment was repeated using a different set of primers, the F8 and R12 forward and reverse primers (Table S1). Highest expression was again observed in the spleen with significant expression observed in the parietal lobe and corpus callosum with lower levels in the cerebral cortex and temporal lobe (Fig. S2b, lanes 1, 8, 9, 4 and 7 respectively). With both sets of primers, namely the [F7 and R10] and [F8 and R12] pairs, the sizes of the observed RT-PCR products correlated precisely to those expected for expression of *LY6S* [Fig. 3di, di’ and dii, and Fig. S2a-for the [F7 and R10] primer pair; Fig. 3f and g (see below), and Fig. S2b and 3c for the [F8 and R12] primer pair].

Whereas RT-PCR analyses revealed significant *LY6S* expression in human spleen tissue, similar analyses performed on a number of different human cell lines failed to detect *LY6S* expression. Both *Ly6a*, as well as other mouse Ly6 genes such as *Ly6c1* (immediately distal to *Ly6a*) are known to be highly induced by interferons and human *LY6E* located immediately upstream to *LY6S* is induced by both Type I (alpha/beta) and Type II (gamma) interferons. We thus postulated that *LY6S* may also be interferon-inducible. A perfect interferon-stimulated response element (ISRE) positioned about 500 bp upstream to the *LY6S* gene supports this notion. The ISRE appears as *TTCCTGT<u>GAA</u>*<u>ATGGAAA</u>TTCAGGA (underlined sequence is identical to the ISRE conferring interferon inducibility on one of the human IFN-alpha genes), and conforms precisely to the tandem GAAANNGAAA element that appears in most Type I Interferon Stimulated Genes (ISGs). Furthermore, this same stretch of 24 nucleotides contains the small palindromic consensus sequence *<u>TTC</u>*N2-4*<u>GAA</u>* (*<u>TTC</u>CTGT<u>GAA</u>*) that conforms to the consensus gamma interferon activation site (GAS). We thus assessed *LY6S* expression in different human cell lines, either untreated or exposed to IFN-alpha, IFN-beta or IFN-gamma, and compared *LY6S* expression with that of *LY6E*. Because of the number of PCR cycles used in this experiment (36 cycles), *LY6E* expression was observed at high levels in all cell lines irrespective of IFN treatment (Fig. 3f, upper panel). In contrast, in the absence of cytokine treatment none of the human cell lines expressed significant levels of *LY6S*, as shown for the M12CB3 and YCB3 melanoma cells (Fig. 3g, lane 1, and Fig. S2c, panels i and ii, lanes 1), and for the MCF7 breast cancer cells (Fig. S2c, panel iii, lane 1). Treatment with IFN-beta induced marked *LY6S* expression in several cell lines including U87 glioblastoma cells (Fig. 3f, lanes 8 and 12), MCF7 breast cancer cells (Fig. 3f, lane 18 and Fig. S2c, panel iii, lane 2) and HEY ovarian cancer cells (Fig. 3f, lane 17). Interferon treatment induced LY6S expression in M12CB3 melanoma cells (Fig. 3g, lanes 2, 3 and 4, for interferons alpha, beta and gamma, respectively and Fig. S2c, panel i, lane 2) and in YCB3 melanoma cells (Fig. S2c, panel ii, lane 2). Notably, LY6S isoforms 1 and 2 (see below for description of isoforms 1 and 2), were induced to varying extents by the different interferons (Fig. 3g, compare lanes 2, 3 and 4).

TOPO cloning and nucleotide sequencing of the *LY6S* RT-PCR product(s) obtained with human spleen cDNA (see Fig 3d-i, lane 18) revealed three inferred protein isoforms (Fig. 4 and Fig. S3 for detailed information on LY6S-iso1 and iso2) corresponding to: (1) a LY6S protein, which received a perfect GPI score from the GPI anchor predictor tool PredGPI, (Fig. S4) indicating that it is a GPI-linked membrane bound protein (isoform #1, designated LY6S-iso1, in 3 out of 8 sequenced TOPO clones, Fig. 4a, isoform #1), (2) a C-terminally truncated LY6S protein, likely to be secreted from the cell that retains both exon 1 (SP) and exon 2 coding sequences, but uses an alternative splice donor site towards the 3’-end of exon 2 (isoform #2, designated LY6S-iso2, for 4 out of 8 sequenced TOPO clones, Fig 4. b, isoform #2) and (3) a protein that retains the exon 1 signal peptide (SP), but by use of an alternative splice acceptor site produces a frame-shifted protein C-terminal to the SP, that is also likely to be secreted from the cell (isoform #3, for 1 out of 8 sequenced TOPO clones, Fig. 4c). All the isoforms conformed to consensus splice donor and splice acceptor sites (*<u>“gt”</u>* and *<u>“ag”</u>* and their flanking sequences, Fig. 4d). Providing additional confirmation, TOPO cloning and sequencing of the RT-PCR products obtained from the interferon treated human melanoma cells (see Fig. 3g) revealed the same LY6S isoforms to those observed in spleen.

**Fig. 4.**
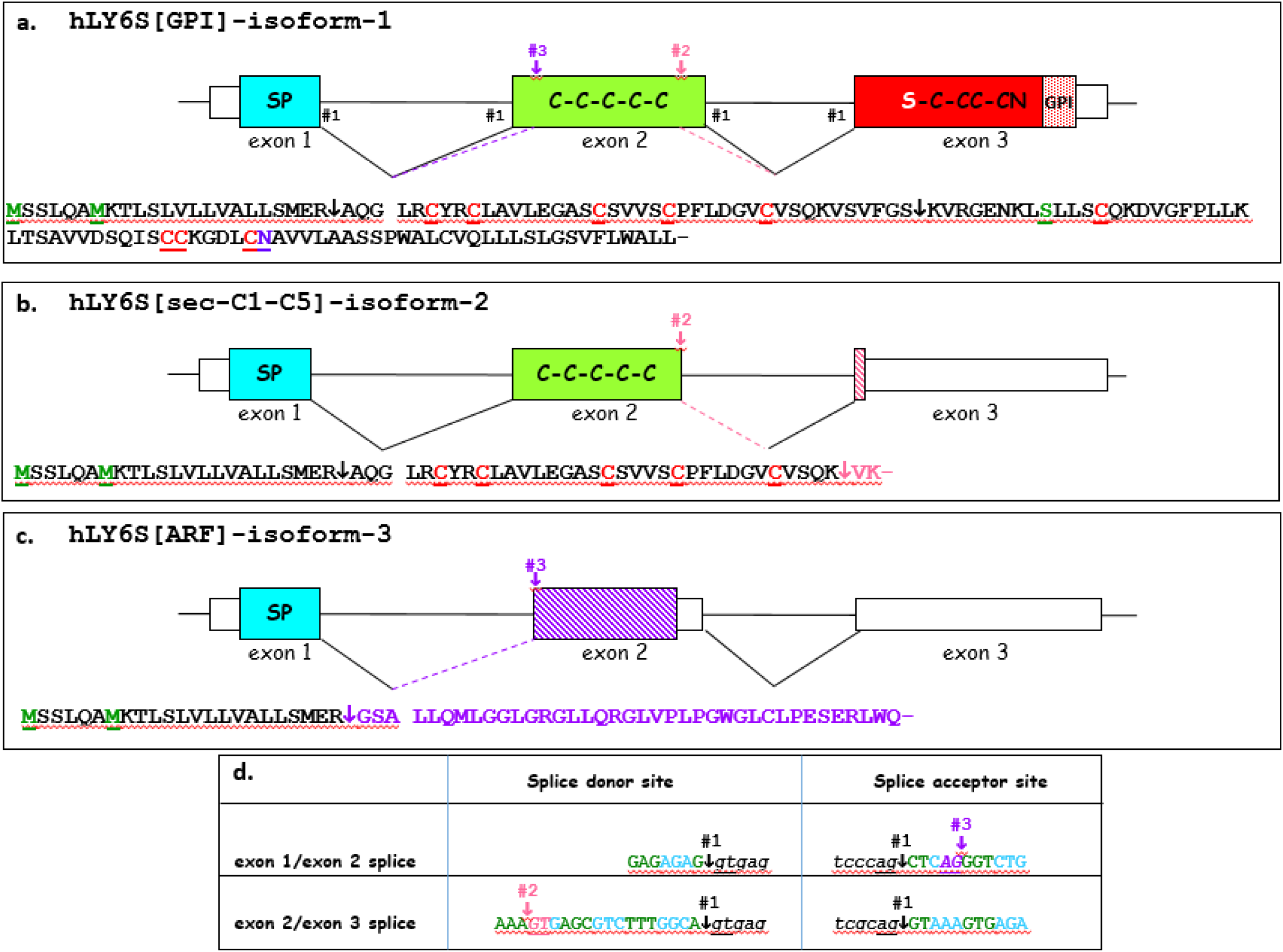
LY6S isoforms determined by sequencing TOPO cloned LY6S spleen RT-PCR products. RT-PCR products obtained from spleen cDNA by using *LY6S* specific forward and reverse primers were TOPO cloned and the inserts were sequenced. The amino acid sequences conceptually derived from the nucleotide sequences are shown for each isoform-LY6-like GPI-linked protein isoform 1, LY6-like secreted protein isoform 2 and secreted protein isoform 3 (panels a, b and c, respectively). The location of splice sites generating the 3 LY6S isoforms was determined by comparing the cDNA nucleotide sequences with the genomic locus of *LY6S*, and are indicated by the hashtag # (panels a, b, c and d). The downward facing arrows in the amino acid sequences designate the corresponding splice sites. The predicted cleavage site of the signal peptide is designated by the gap in the protein sequences. The exon/intron boundary sequences are shown in panel d.

Although belonging to the LY6/uPAR family of proteins which normally contain 10 consensus cysteine residues, the LY6S-iso1 protein described here is special in that it contains a non-canonical serine residue in place of the sixth cysteine residue present in all LY6-like proteins described to date, hence the LY6**<u>S</u>** designation. The serine codon was observed in cDNAs generated from both spleen and interferon-treated melanoma cells and in accord with our findings, all databases investigated show a codon at this position in the *LY6S* gene, in the genomic DNA, coding for a serine residue.

Independent validation for the existence of mRNAs coding for LY6S-iso1 and LY6S-iso2 proteins was provided by analyses of public domain human spleen RNA-Seq data (Fig. S5a, b and c) that showed transcripts derived from the *LY6S* gene in the human spleen.

### LY6S is syntenic to 8 murine Ly6 genes that form the Ly6a subfamily cluster of genes

In contrast to the sole human *LY6S* gene present in the region bordered by *LY6E* and *LY6L*, the syntenic mouse locus within which *Ly6a* resides contains 8 Ly6 genes, all of which are highly similar to each other (Fig. 1a and Fig. 5). These mouse Ly6 genes demonstrate coordinated transcriptional responses to certain inflammatory cytokines, and to those relating to interferon-mediated immune sensing. For example, both *Ly6a* and *Ly6c* are upregulated more than 100-fold in response to a lack of ADAR1 RNA editing activity, the consequence of which is increased sensing of endogenous dsRNA by MDA5 leading to elevated interferon levels, and activation of the innate immune sensing system (*Fig. 3 in (20)*). Additionally expression in intestinal epithelial cells of *Ly6i*, *Ly6a*, *Ly6f* and *Ly6c* in mouse models of inflammatory bowel disease is markedly increased, and inflammatory cytokines including IFN-gamma, TNF-alpha and IL-22 all lead to increased expression of these *Ly6* genes (*21*).

**Fig. 5.**
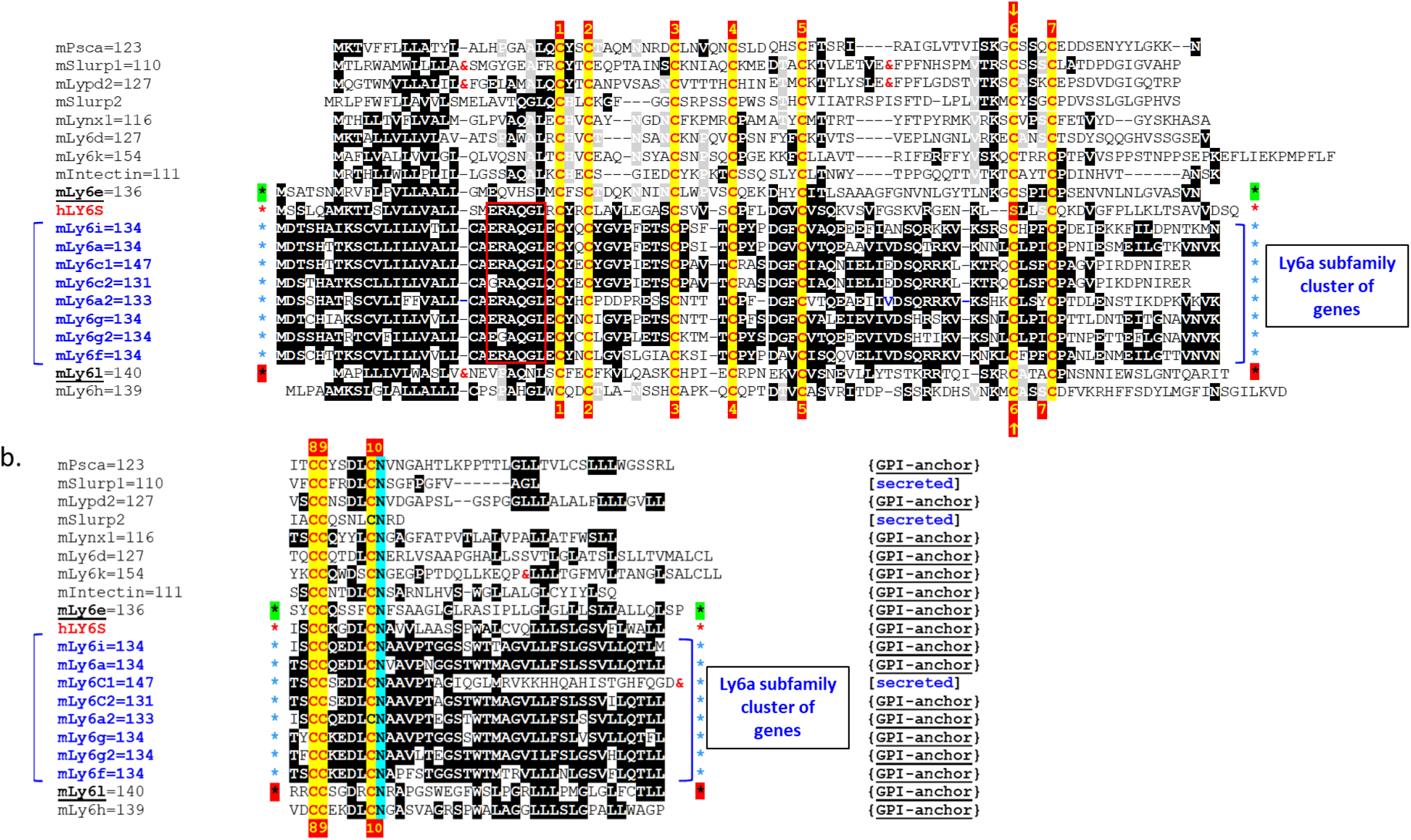
Sequence homology analysis of human LY6S-iso1 together with the Ly6 proteins derived from murine chromosome 15 Ly6 genes. The amino acid sequence of LY6S-iso1 (indicated by a red asterisk) is compared with the sequences of the ‘mChr15 Ly6’ proteins. The Ly6a subfamily genes are bracketed in blue and indicated by the blue asterisks. Identical amino acids appearing 4 or more times in the different Ly6/LY6 proteins are highlighted using a white bold font against a black background. These identities appear at least twice within the Ly6a subfamily cluster proteins and identical amino acids appearing 4 or more times outside of the and not appearing within it are highlighted in bold white fonts against a grey background. The highly conserved amino acid sequence ‘ERAQGL’ appearing in the signal peptides of both the murine Ly6a subfamily cluster proteins and in LY6S-iso1 is boxed by the red rectangle. The Ly6/LY6 consensus cysteine residues of the Ly6 proteins are numbered from 1 -10 above the sequences and indicated by yellow-highlighted red-bold fonts. The unique LY6S-iso1 serine (S) residue appearing in place of the consensus sixth Ly6/LY6 cysteine residue (arrowed yellow 6 against a red background) is indicated by the red highlighted yellow-bold ‘S’ font. The mouse *Ly6e* and *Ly6l* genes bordering the Ly6a subfamily cluster are indicated by black asterisks against green and red backgrounds respectively. Amino acid sequences deleted from the alignment are indicated by the red ‘&’ font. Because of their divergence from other mChr15-Ly6 proteins, Gml, Gml2, Gpihbp1 and mD730001G18Rik=87 were not included in this analysis.

The 8 Ly6 family genes are defined as belonging to the Ly6a subfamily (Fig. 5). Both phylogenetic analyses (Fig. 2) as well as protein similarity analyses (Fig. 5) indicate that of all the mChr15-Ly6 proteins, the human LY6S-iso1 protein is most similar to the proteins encoded by the Ly6a subfamily. All murine genes within this subfamily code for proteins containing the amino acid sequence ‘ERAQGL’ within the C-terminal regions of their signal peptide. This sequence is also conserved in the human LY6S protein (boxed red rectangles in Fig. 5 and Fig. S6a), yet it is not present in any of the other human chr8 LY6 proteins.

The constructed phylogenetic maximum likelihood tree (Fig. 2) shows *LY6S* branching at the base of the Ly6a subfamily clade containing the 8 *Ly6* mouse genes located within the 500kb sub-region. A bootstrap value of 89% shows that the phylogenetic signal from the whole alignment lends strong support to this grouping. This tree supports the notion that *LY6S* has evolved from the same ancestral gene as the *Ly6a* subfamily clade and that there have been several duplication events in the mouse. The similarity of *LY6S* to the 8 *Ly6a* subfamily genes as well as their syntenic colocalization (both flanked by *LY6E/Ly6e* and *LY6L/Ly6l*), indicates that the 8 Ly6a subfamily genes are homologous to the solitary human *LY6S* gene and this is a 1:8 orthology relationship (Fig. 1, Fig. 2 and Fig. 5). Comparison with other well annotated vertebrate genomes suggest that the most parsimonious explanation is that there has been a gene expansion of *Ly6* genes in mouse, rather than one or more deletion events in other species.

*LY6S* has not been annotated in other species, and its absence from human gene annotation may have biased automatic annotations of other vertebrate genomes against discovery of *LY6S* orthologs. Furthermore, the annotation of *LY6S* orthologs in primate genomes containing an ortholog of *C8orf31* may be hindered in a manner similar to that in humans. BLAT searches using the LY6S protein sequence as a query suggest that some other primates are likely to have copies of the *LY6S* gene. Although the *LY6E* to *LY6L* region is missing from the latest chimp assembly (2018 Clint_PTRv2/panTro6), it does appear in the previous assembly (2016 Pan_tro 3.0/panTro5).

A protein BLAST search with LY6S-iso1 yielded as the best hits the lymphocyte antigen Ly6a (Ly6A-2/Ly6E-1) proteins from Nannospalax galili (Upper Galilee Mountain blind mole rat), Microcebus murinus (grey mouse lemur), Merionus unguiculatus (Mongolian gerbil) and Peromyscus maniculatus bairdii (North American deer mouse) with e-values of 1e-38, 3e-30, 2e-28 and 7e-28, respectively [see Fig. S6a for a comparison of LY6S-iso1 with the Nannospalax and Peromyscus Ly6a (Ly6A-2/Ly6E-1) proteins]. Subsequent pairwise comparison of each human LY6 protein present on chromosome 8 with the deer mouse Ly6A-2/6E-1-like protein showed that LY6S gave the best match by far (Fig. S6c, d and e).

### LY6-iso1 protein is expressed by interferon-treated cells and by spleen cells

To facilitate investigation into *LY6S* expression at the protein level, LY6S-iso1-specific monoclonal antibodies designated 5E12 were generated, the sequence of which is presented (Fig. S7). The specificity of 5E12 was confirmed by the following: (1) 5E12 bound to recombinant hFc-LY6S fusion protein (Fig. 6b, lane 6) and binding was blocked by the addition of the immunizing LY6S peptide-E and a peptide contained within this sequence (Fig. 6b, lanes 7 and 9, respectively, and see Fig. 6a for peptide sequences within the LY6S-iso1 protein) but not by peptide C, which is upstream to the immunizing peptide (Fig. 6b, lane 8), (2) 5E12 bound to a protein expected for the size of the LY6S-iso1 protein in western blot analyses of HK293 cells infected with lentiviruses coding for the LY6S-iso1 protein (Fig. 6c, left panel lanes 1 and 2), but not from cells infected with control lentiviruses (Fig. 6c, left panel lane 3). Furthermore, LY6S-iso1 protein observed in cells infected with retroviral particles coding for the native LY6S-iso1 protein migrated slightly faster than the corresponding LY6S protein that contained a Flag-epitope at its N-terminus, conforming precisely to their difference in molecular mass [compare in Fig. 6, panel c, lane 1 (with the Flag epitope at the N-terminus of the mature protein) and lane 2 (LY6S protein without the Flag epitope)]. (3) By flow cytometry, 5E12 detected cell-surface LY6S-iso1 protein, albeit at low levels, on intact HK293 cells infected with retroviral particles coding for LY6S-iso1, as expected for a GPI-linked protein (Fig. 6d, Panels ii and iii), and (4) anti-LY6S mAb 5E12 immunofluorescent analysis of HK cells transiently transfected with plasmids coding for the LY6S protein showed positively staining cells (Fig. 6e, panels ii and iii), that were not seen with cells transfected with control empty plasmid (Fig. 6e, panel i). The reproducible detection of expression at the cell surface at low levels, (Fig. 6d, panels ii and iii) suggests that cells do not readily express the LY6S protein at the cell surface (and see below).

**Fig. 6.**
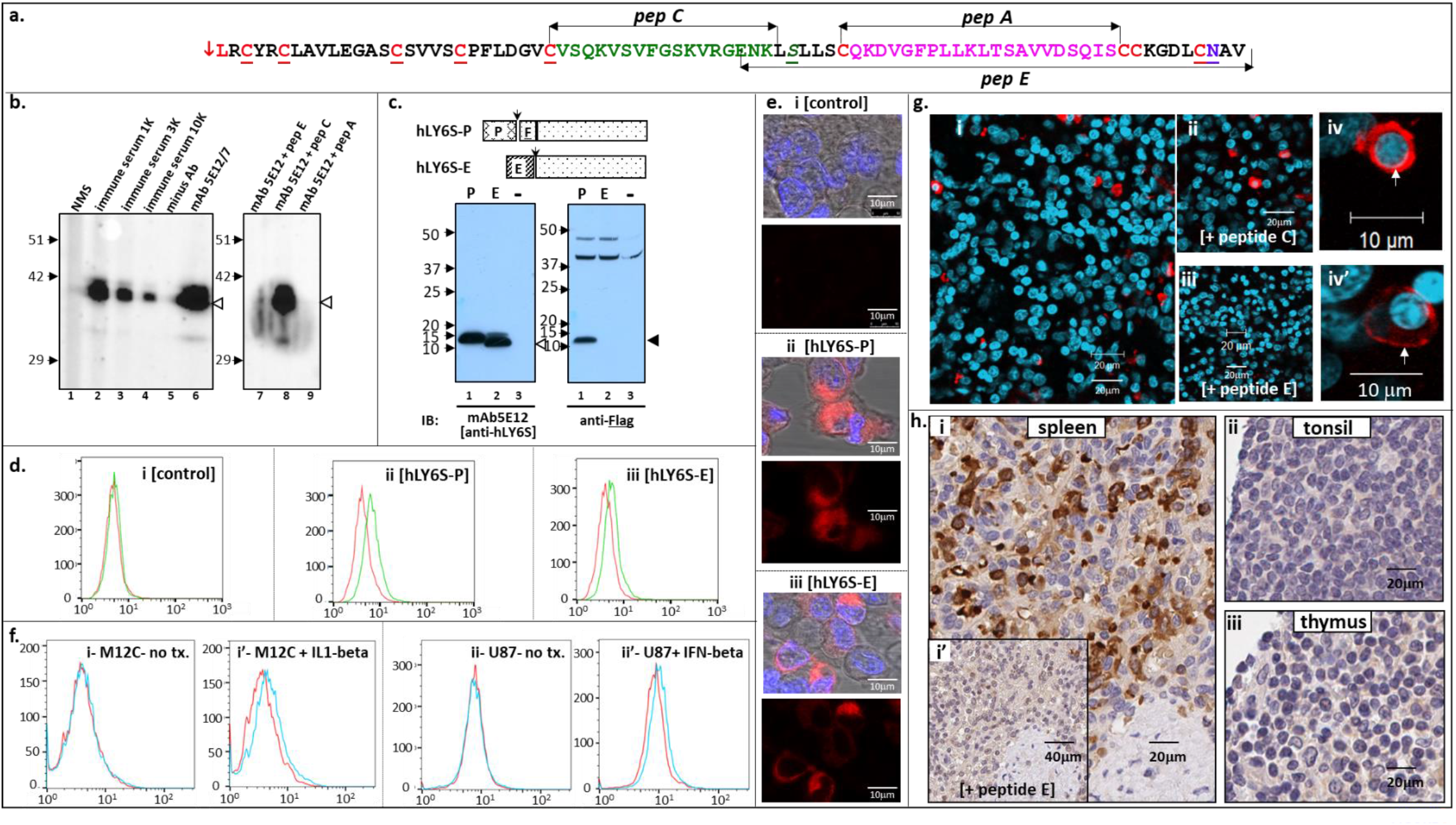
Detection of LY6S-iso1 protein with anti-LY6S-iso1 mAb 5E12 on the cell surface and in human spleen tissue. **Panel a-** Peptides A, C and E, as indicated derived from the LY6S-iso1 protein sequence were synthesized and mice were immunized with Peptide E. **Panel b-** The anti-LY6S-iso1 polyclonal antibodies (serum dilutions indicated) from a peptide-E immunized mouse or anti-LY6S-iso1 mAb-5E12 generated from the spleen of this mouse were used to probe an immunoblot of SDS-PAGE resolved hFc-LY6S-iso1 fusion protein (lanes 2-4, and lane 6 respectively)-normal mouse serum served as control (lane 1). Similarly, an immunoblot of hFc-LY6S-iso1 protein was probed with anti-LY6S-iso1 mAb-5E12 in the presence of peptides E, C and A (lanes 7-9, respectively). **Panel c-** HEK293 cells were stably transfected with expression plasmids coding for the native LY6S-iso1 protein (hLY6S-<u>E</u>, lanes indicated by E) or for hLY6-P in which the <u>E</u>ndogenous signal peptide (SP) was replaced by the <u>P</u>reprotrypsin SP followed by the Flag epitope (hLY6S-<u>P</u>, lanes indicated by P). Immunoblots of cell lysates prepared from these stably transfected cells (P, E or control [-], lanes 1, 2 and 3 respectively) were probed with mAb-5E12 (anti-LY6S-iso1) and with anti-Flag (left and right panels, respectively). **Panel d-** Non-transfected HEK293 cells and cells stably transfected with expression plasmids coding for LY6S-P or LY6S-E (panels i, ii and iii, respectively) were investigated by flow cytometry with anti-mouse secondary antibody alone (red lines) or with mAb-5E12 followed by fluorescently labelled secondary antibodies (green lines). **Panel e-** Non-transfected and HEK293 cells stably transfected with LY6S-P or LY6S-E (panels i, ii and iii, respectively) were stained immunofluorescently with mAb5E12. **Panel f-** Human melanoma (M12C, panels i and i’) or human glioblastoma (U87, panels ii and ii’) cells were grown in the absence of cyotokines (panels i and ii) or with IL1-beta (panel i’) or interferon-beta (panel ii’). The cells were assessed by flow cytometry with secondary antibody alone (red lines) or with mAb-5E12 followed by the fluorescently labelled anti-mouse secondary antibodies (green lines). **Panel g-** Sections of human spleen tissue were stained immunofluorescently with anti-LY6S-iso1 mAb-5E12 in the absence of peptides (panel i), or in the presence of peptide C (panel ii) or peptide E (iii). LY6S-iso1^+^ spleen cells at higher magnification are shown in panels iv and iv’. **Panel h-** Immunohistochemical staining of sections of human spleen, tonsil and thymus with anti-LY6S-iso1 mAb-5E12 (panels i, ii and iii, respectively) and of spleen with anti-LY6S-iso1 mAb-5E12 in the presence of competing peptide E (panel i’). Cells stained brown are LY6S-iso1^+^ cells.

Human M12 melanoma cells treated with IL1-beta (Fig. 6f, panel i’) as well as with interferon-gamma and IL6 (Fig. S8, panels i’’ and i’’’) demonstrated cell-surface LY6S protein expression. Similarly, U87 glioma cells treated with interferon-beta showed cell-surface LY6S protein (Fig. 6f, panel ii’), as did cells treated with IL1-beta (Fig. S8, panel ii’’). Thus, cytokine treatment of cells reproducibly induces the expression of cell surface LY6S in both M12 melanoma and U87 glioma cells. Such expression is not detected in the absence of cytokine treatment (Fig. 6f, panels i and ii).

Immunofluorescent staining of human spleen tissue provided further support for the endogenous expression of the LY6S-iso1 protein in human cells. This showed a discrete sub-population of LY6S-iso1-positive cells that accounted for about 5% of all spleen cells (Fig. 6g, panel i). The immunizing peptide E abrogated staining (Fig. 6g, panel iii) confirming antibody specificity, whereas peptide C which was not used for immunization (Fig. 6g, panel ii) had no effect. LY6S-iso1 localized to the cell surface (Fig. 6g, panel iv’), but was also observed within the cell (Fig. 6g, panel iv). LY6S-iso1 protein expression was barely detectable in other lymphoid tissues such as thymus and tonsils, as assessed by immunohistochemical staining (Fig. 6h, panels ii and iii), yet was seen clearly in spleen cells (Fig. 6h, panel i), stained under identical conditions. Spleen tissue costained for LY6S-iso1 together with antibodies directed against additional cell-surface proteins that define particular cell types, detected a prominent sub-population of LY6S-positive cells constituting about 5% of all the human spleen cells (Fig. 6g-panel i, 6h-panel i, Fig. 7-panels b’ and b’’ and Fig. 7-panels e-p). Staining of thymus, tonsil and spleen, with anti-CD45 a pan-leukocyte marker (Leukocyte Common Antigen, LCA) showed many CD45-positive cells in all three lymphoid-rich tissues (Fig. 7a, b and c and Fig. S9, panels a’, b’, c’ and d’). Costaining with anti-LY6S antibodies revealed that only the spleen contained LY6S-positive cells (Fig. 7b’, Fig. S9, panels b” and d”), distinguishing it from the LY6S-negative thymic and tonsillar tissues (Fig. 7a’, and c’ and Fig. S9, panels a” and c”). Antibodies against CD8 (Fig. 7e), CD4 (Fig. 7f), CD34 (Fig. 7g), CD19 (Fig. 7h), CD11b (Fig. 7i, 7j), FoxP3 (Fig. 7k), CD11c (Fig. 7l), CD20 (Fig. 7m), CD117-cKit (Fig. 7n), CD68 (Fig. 7o) and CD31 (Fig. 7p) showed no colocalization of any of these markers with LY6S-iso1 positive cells, suggesting that the LY6S-positive cells do not belong to a classical lineage of lymphoid cells. In contrast to the other tested leukocyte markers, a discrete subpopulation representing about 10% of the LY6S-positive splenic cells also expressed the Leukocyte Common Antigen (LCA) marker CD45 (Fig. 7, panel b’’, and panels d), indicating that a subset of LY6S-iso1+ve and CD45+ve cells express both proteins.

**Fig. 7.**
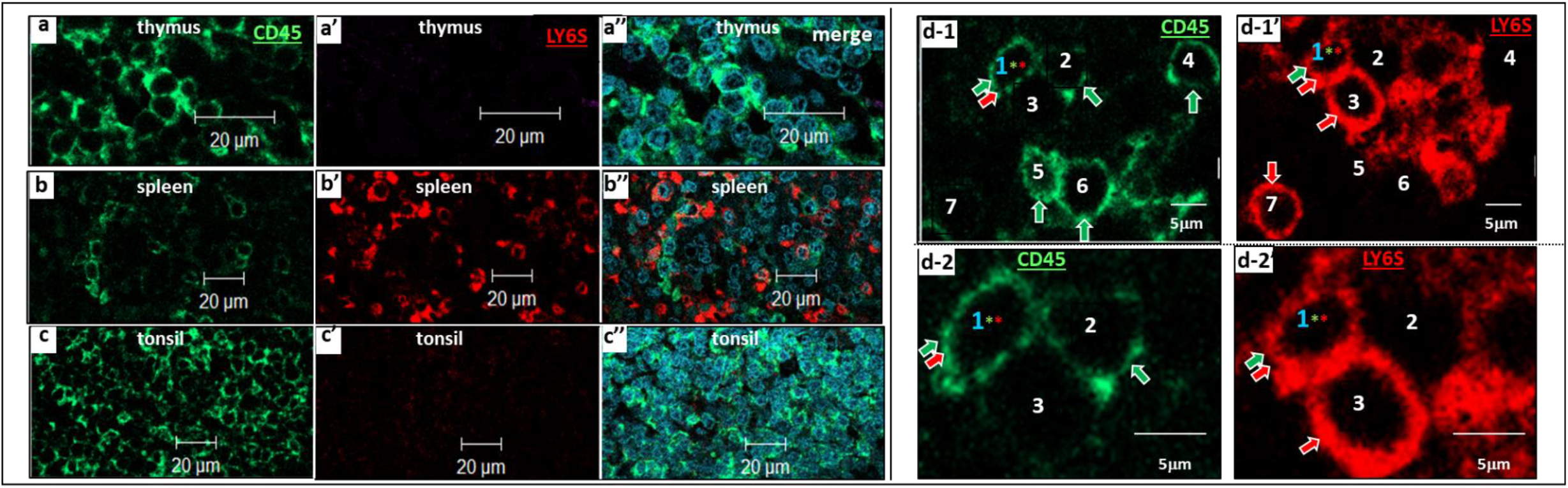
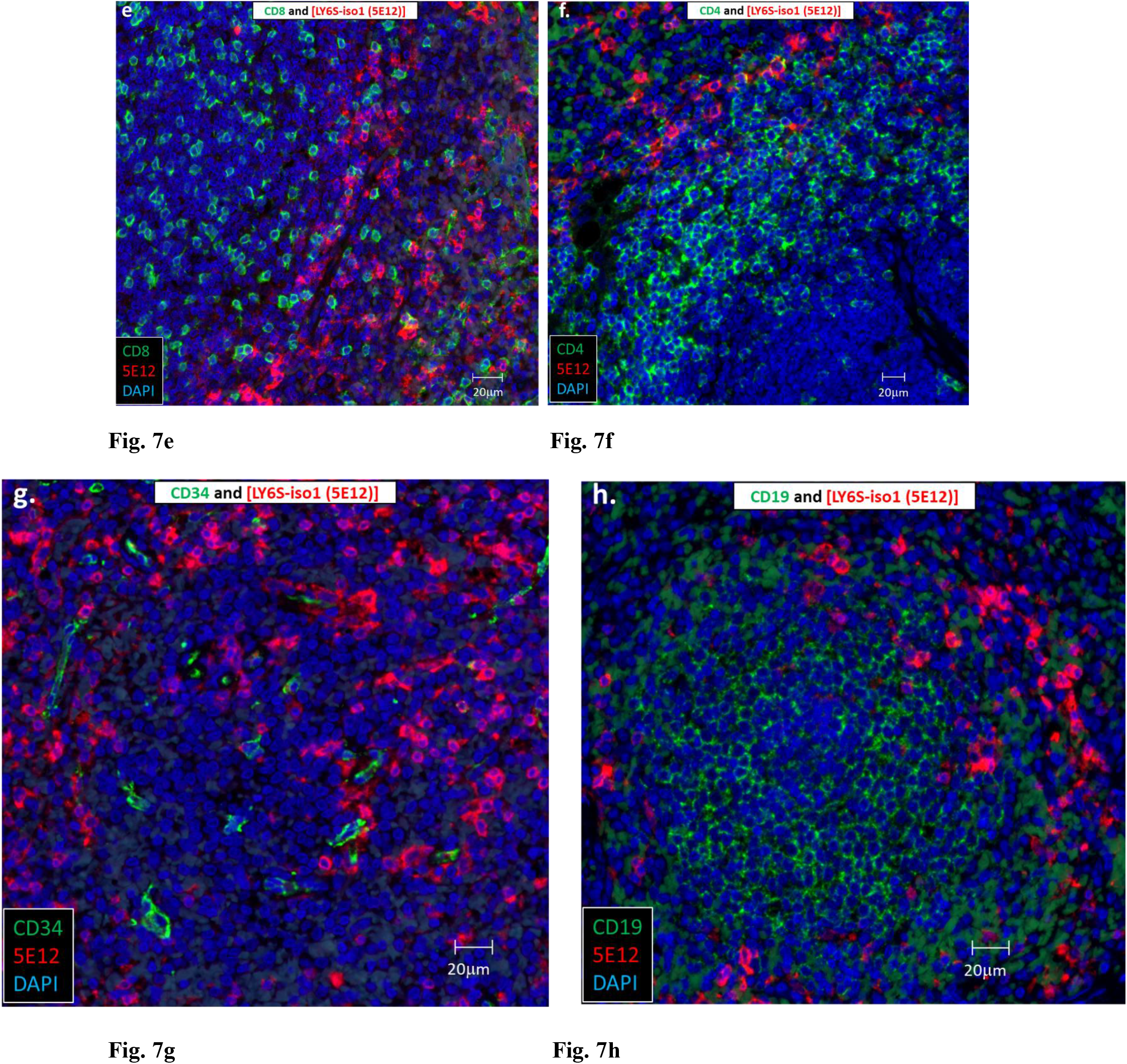
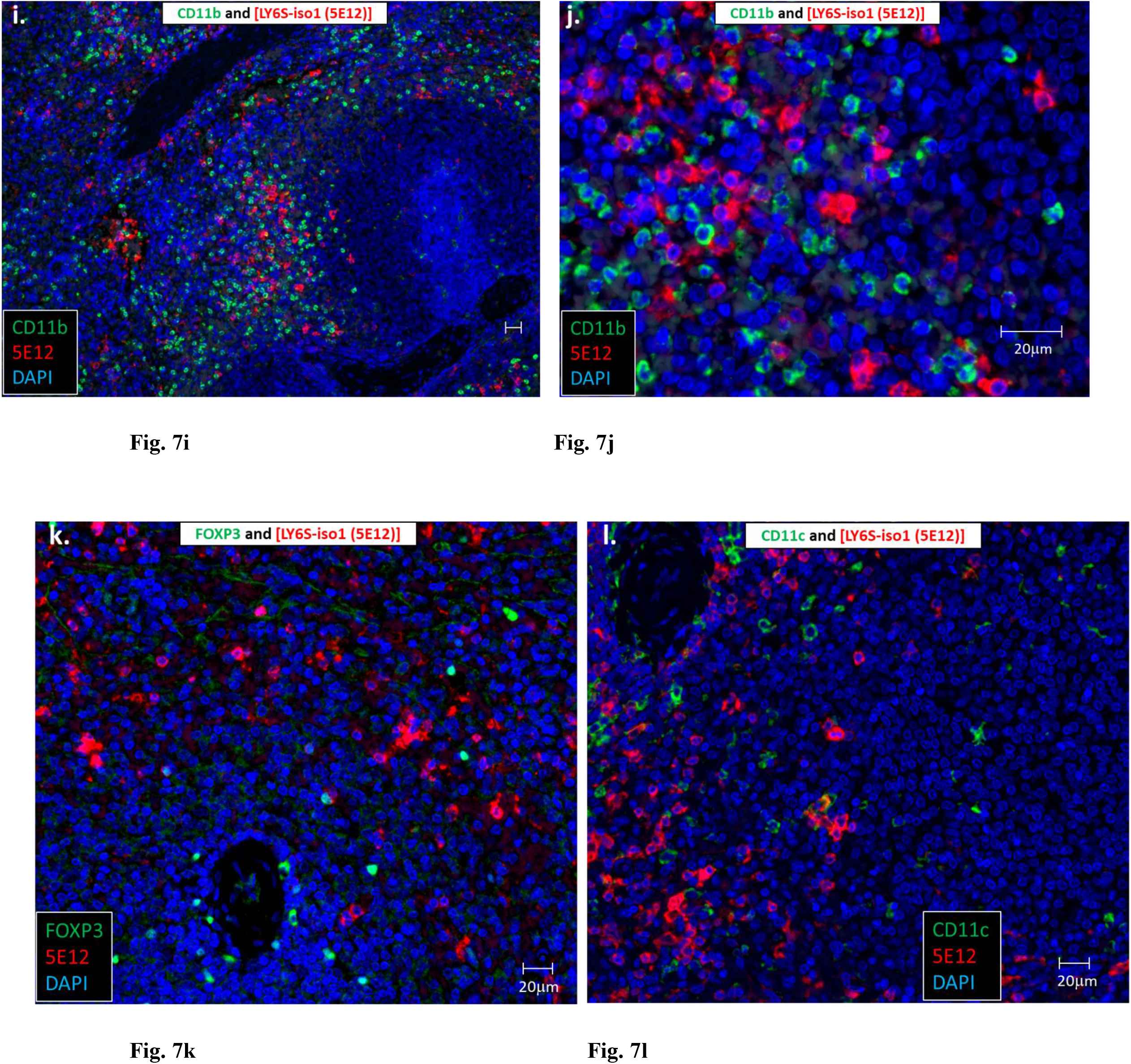
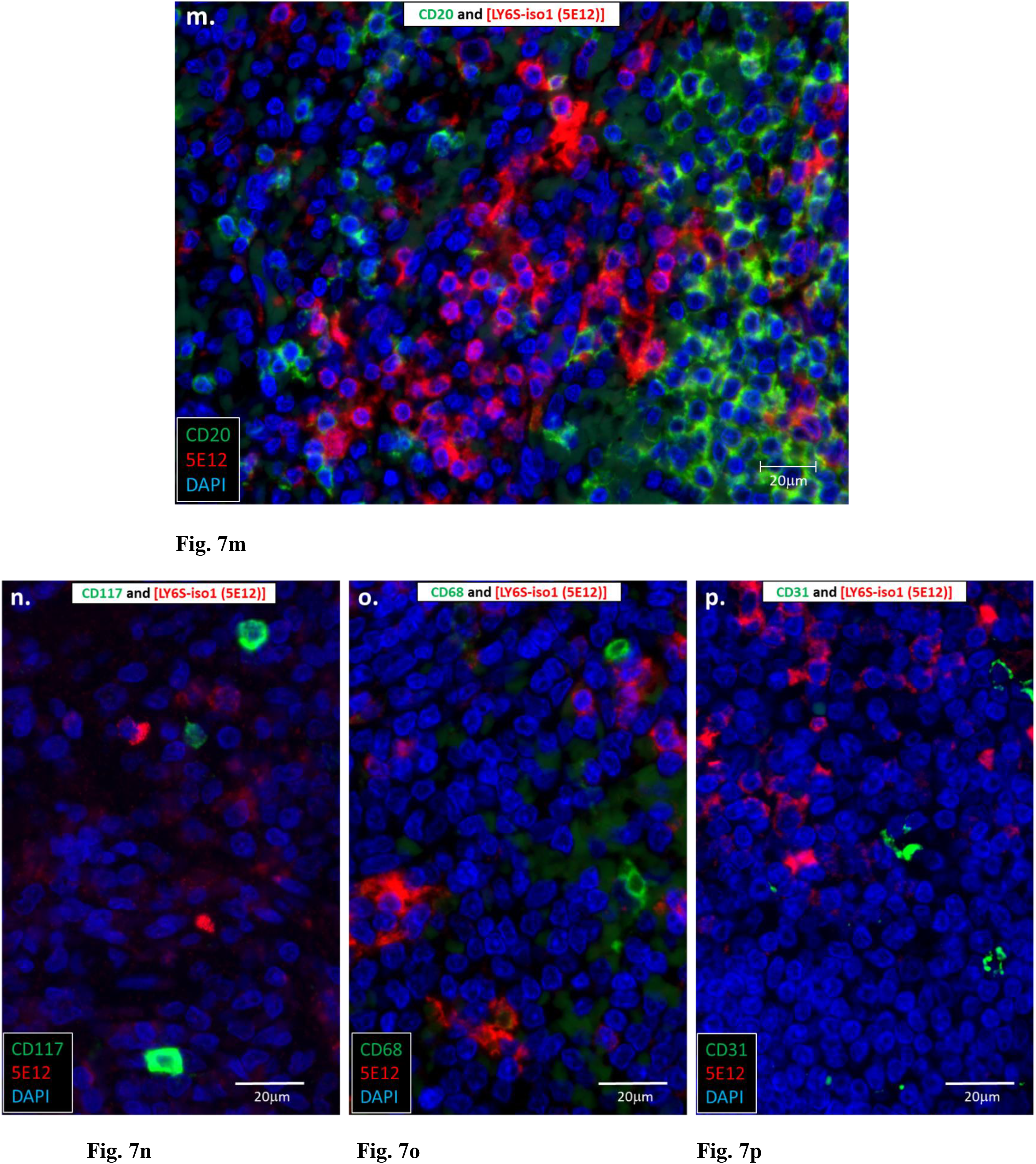
Immunofluorescent staining for the LY6S-iso1 protein and other cell-marker proteins in spleen, thymus, and tonsil. **Panels a-d-** Sections of thymus, spleen and tonsil were co-immunofluorescently stained for CD45 and LY6S-iso1 (green, panels a-c, and red panels a’-c’, respectively). Merging of the two stains is shown in panels a’’, b’’ and c’’. Two different selected fields of spleen (d-1, d-1’ and d-2, d-2’) shown at two higher magnifications were stained immunofluorescently for CD45 (green, d-1 and d-2, CD45^+^ cells indicated by green-filled arrows) and for LY6S-iso1^+^ (red, d-1’ and d-2’, red-filled arrows). The doubly positive CD45^+^/LY6S-iso1^+^ cell in this field is cell number 1. **Panels e-p** (below)-Spleen sections immunofluorescently stained for LY6S-iso1 (in red) and the indicated cell marker proteins (in bright green)- the dull green staining seen in some sections is non-specific background staining).

### Expression of the LY6-iso1 protein elicits changes in cell growth

The findings (a) that LY6S expression levels in adult tissues, likely exposed to foreign pathogens such as bacteria and viruses, were at all times higher than those seen in fetal tissues (Fig. 3e, inset) and (b) that LY6S expression was observed only following cytokine treatment of cells (Fig. 3f and 3g, Fig. 6f, and Fig. S8) suggested that LY6S expression may be related to an activated cell phenotype. To test this supposition, human cell lines were infected with retroviral particles coding for the GPI-linked cell-surface LY6S-iso1 protein or for the secreted LY6S-iso2 protein. Cells expressing the secreted LY6S-iso2 protein showed no differences in gross morphology, compared with control cells stably infected with control pQCXIP plasmid (Fig. 8a-i, compare right inset [LY6-iso2] with left inset [control vector]). However cells infected with particles coding for the cell-surface linked LY6S-iso1 protein showed changes in cell morphology (compare cells expressing LY6-iso1 in Fig. 8 panels a-ii, a-ii’, b-ii, b-ii’ and c-ii, c-ii’ with cells expressing control vector in Fig. 8 panels a-i, a-i’, b-i, b-i’ and c-i, c-i’), where cells expressing LY6S-iso1 protein were large and many contained prominent cytoplasmic vacuoles (see for example Fig. 8, panel a-ii’). Cells displaying this phenotype included U87 (glioma, Fig. 8a-ii and a-ii’), MCF7 (breast carcinoma, Fig. 8b-ii and b-ii’) and YDFR (melanoma, Fig. 8c-ii and c-ii’). The LY6S-iso1-expressing puromycin-resistant cells initially grew slowly (as shown for YDFR melanoma cells expressing LY6S-iso1, Supplementary Data, Fig. 11). However, after three to four passages, the LY6S-iso1 cells resumed growth, reached plate confluency and could then be subcultured.

**Fig. 8.**
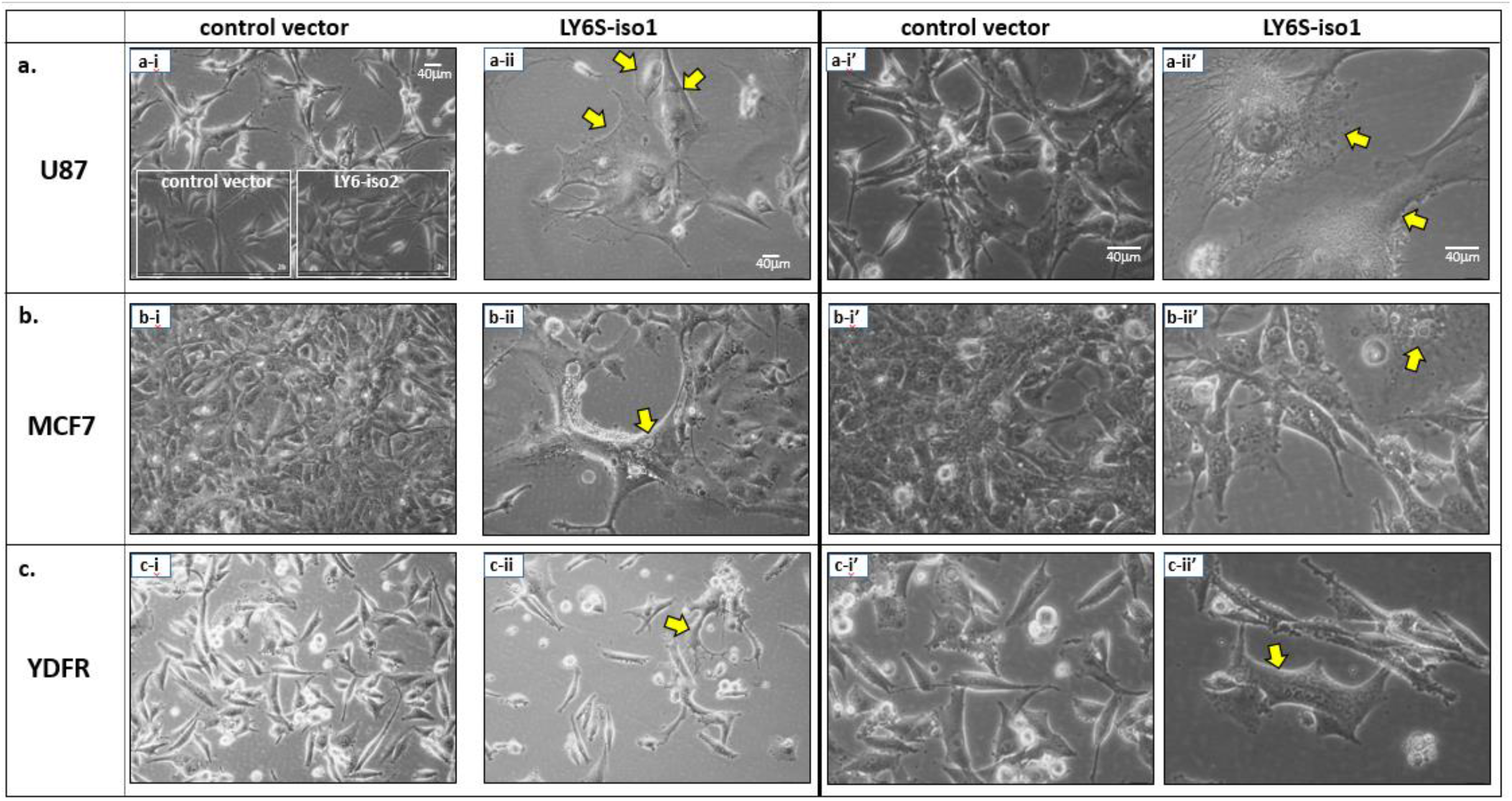
Cell morphology of human U87 glioblastoma, MCF7 breast carcinoma and YDFR melanoma cells transfectants expressing LY6S proteins. U87, MCF7 and YDFR cells (panels a, b and c, respectively) stably transfected with pQCXIP-control vector (panels a-i, b-i, c-i and a– i’, b-i’, c-i’) or with pQCXIP coding for LY6S-iso1 protein (panels a-ii, b-ii, c-ii and a–ii’, b-ii’, c-ii’) were photomicrographed at low and high magnification (left and right sets, respectively). Exceptionally large cells with prominent vacuoles are indicated by yellow arrows. A comparison of control U87 cells with U87 cells expressing LY6-iso2 is shown in the 2 panel a-i insets.

### Cells expressing the LY6S-iso1 protein have increased resistance to viral infection

Because (a) transcription of the *LY6S* gene is interferon inducible and (b) LY6S-iso1 expression is associated with changes in cell morphology and growth, we investigated whether expression of the LY6S-iso1 protein impacts cell resistance to viral infection. In all three human cell lines investigated, M12, YDFR and U87, LY6S-iso1 protein expression was associated with a marked diminishment in vesicular stomatitis virus (VSV) viral replication (Fig. 9). In M12 melanoma cells, LY6-iso1 expression led to a decrease in viral yield that exceeded two orders of magnitude (Fig. 9, panels c and d), and a marked decline of viral yields was also seen in human YDFR melanoma and human U87 glioblastoma cells that express the LY6-iso1 protein (Fig. 9, panels a, b and d).

**Fig. 9.**
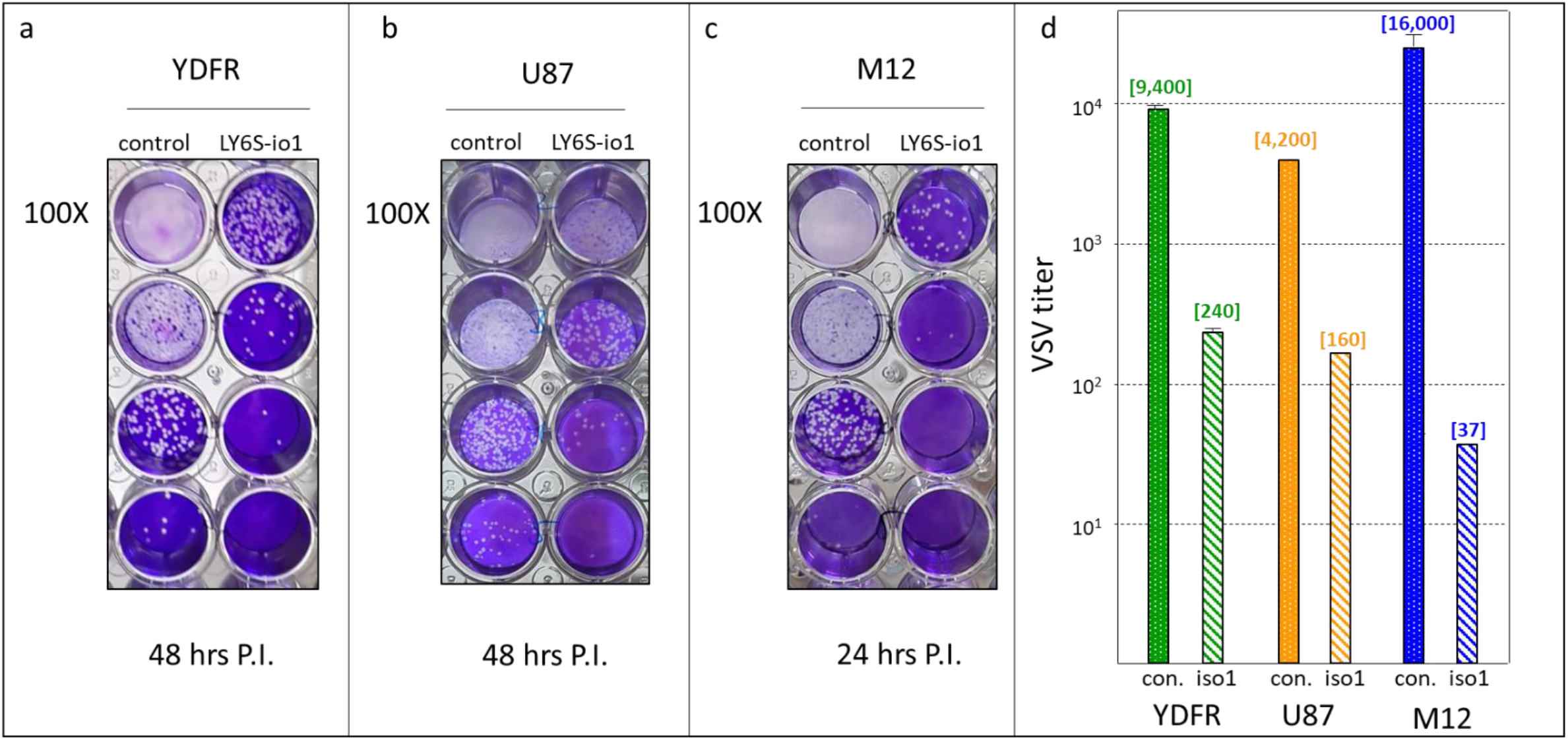
Inhibition of viral replication in cells expressing the LY6S-iso1 protein. Human melanoma cells (YDFR-CB3 and M12-CB3) or human glioblastoma cells (U87), as indicated, were stably transfected with an empty expression vector (control) or with an expression vector coding for LY6S-iso1 (LY6S-iso1). VSV was added to the cell cultures and virus present in the spent medium was assayed on monkey Vero cells, starting with a 100-fold dilution, followed by ten-fold dilutions. Quantitation of the viral titer in the spent medium of the virally infected cultures is shown in panel [d] (for YDFR cells- at 100-fold dilution, 48 hr. time point and MOI of 0.05, for U87 cells- at 1000-fold dilution, 48 hour time point and MOI of 0.015 and for M12 cells- at 100-fold dilution, 24 hr. time point and MOI of 0.005). The number of viral plaques is indicated in square brackets above the bars, for the respective cell types.

### Expression of the LY6-iso1 protein elicits an inflammatory cell phenotype

The changes observed in the phenotype of LY6S-iso1 expressing cells, as described above, suggested changes in the gene expression profiles of these cells. To test for this, next generation sequencing (both Illumina single read sequencing and Paired End (PE) True Stranded) was performed on RNA isolated from triplicate U87 glioblastoma cell cultures infected with control pQCXIP plasmid (U87-control) or from cells stably expressing cell-surface linked LY6S-iso1 protein (designated ‘U87-LY6S-iso1’). Genes coding for chemokines, chemokine receptors, cytokines and other proteins associated with an inflammatory cell phenotype were markedly upregulated in the LY6S-iso1 expressing cells [Fig. 10, shown as volcano (at left) and heat map (at right) plots, and see Table S2]. These included the following: ***Cytokines*:** IL1A, IL1B, IL6, IL11, IL24, IL36B, CD274 (PDL1), CSF2, CSF3, EREG, FGF2, GDNF, HBEGF, LIF and TNFSF8; ***Chemokines*-** CXCL1, CXCL2, CXCL3, CXCL5, CXCL8, CX3CL1 and CCL20; ***Chemokine Receptors*:** ACKR3, CMKLR1 and IL36RN; ***Genes Directly Related to Inflammation and Immune Responses:*** BCL2A1, BGN, LCN2, LCP1, NFkB1, NLRP12, RCAN1, PTGS2, RGS16 and SDC3; (*downregulated*)- CXADR, DEPTOR, DKK1, HMOX1; ***Related to Interferon and Cytokine Signaling:*** IRF5, NFKBIZ and SOCS1; (*downregulated*)- ID1 and SNAI1. Ingenuity analyses (Qiagen IPA as detailed in Methods) of the RNA-Seq data identified the following biological pathways with very high p-values-Canonical Pathways [-log(p-value)] and Annotations of “Diseases and Functions”(p-value): (a) Granulocyte adhesion and diapedesis [12.8], (b) Role of cytokines in mediating communication between cells [9.86], (c) Differential regulation of cytokine production in macrophages and T-helper cells by IL-17A and IL17-F [6.89], (d) Role of pattern recognition receptors in recognition of bacteria and viruses [6.82], (e) Acute inflammatory response (2.99E-13), (f) Accumulation of myeloid cells (1.26E-17), (g)

**Fig. 10.**
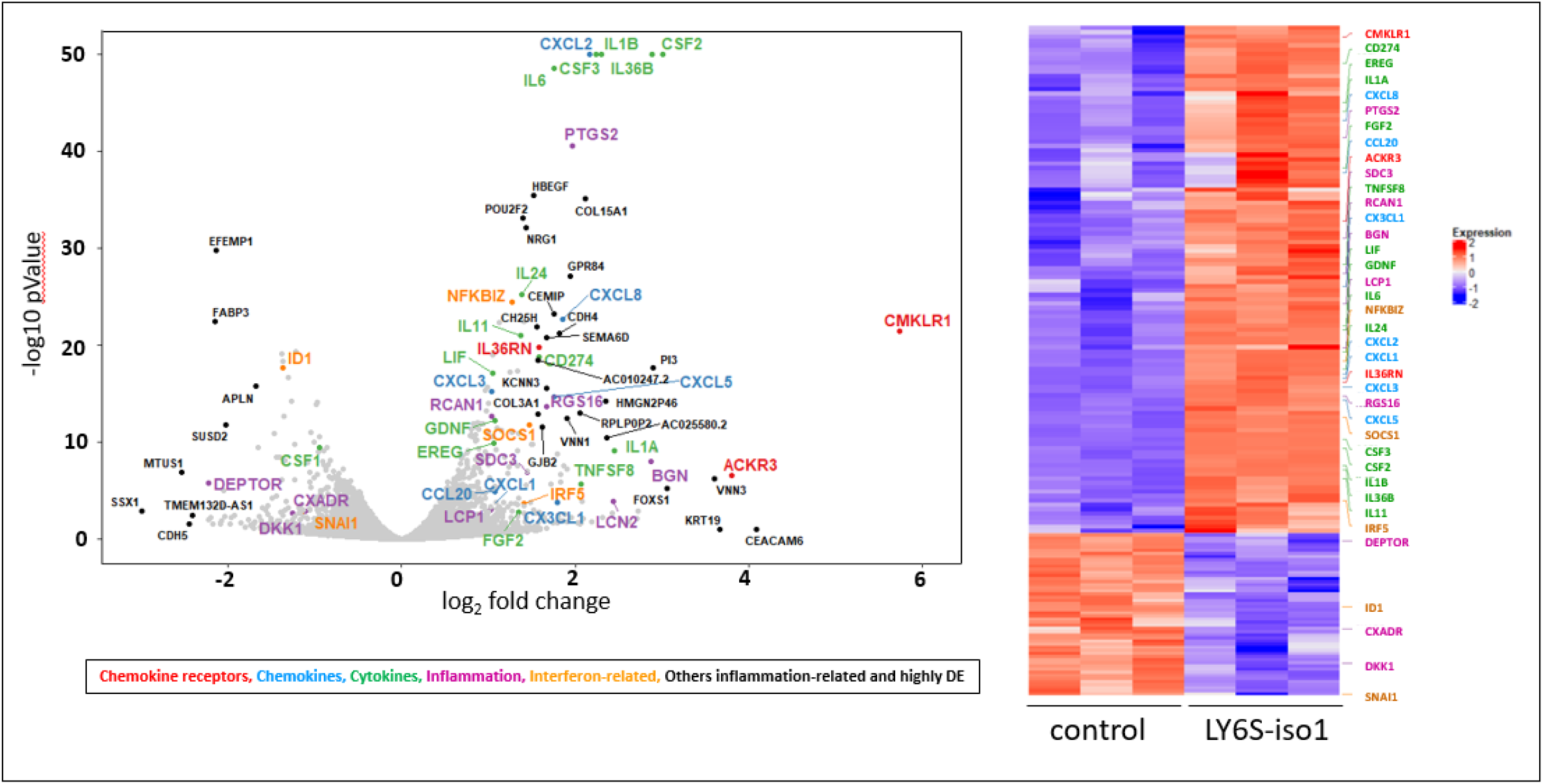
Volcano plot and heat map of genes differentially expressed in LY6S-iso1-expressing U87 cells. RNA extracted from triplicate cultures of puromycin-resistant U87 cells stably transfected with pQCXIP vector coding for LY6S-iso1 or with vector alone were subjected to RNA-Seq analysis as described in Methods, and the differential gene expression between the two groups is represented as a volcano plot and as a heat map (at left and right, respectively). A selection of the differentially expressed genes was color coded as follows: chemokines-light blue, chemokine receptors-red, cytokines-green, inflammation-purple, interferon related-orange, others related to inflammation and also differentially expressed to a great extent-black.

**Fig. 11.**
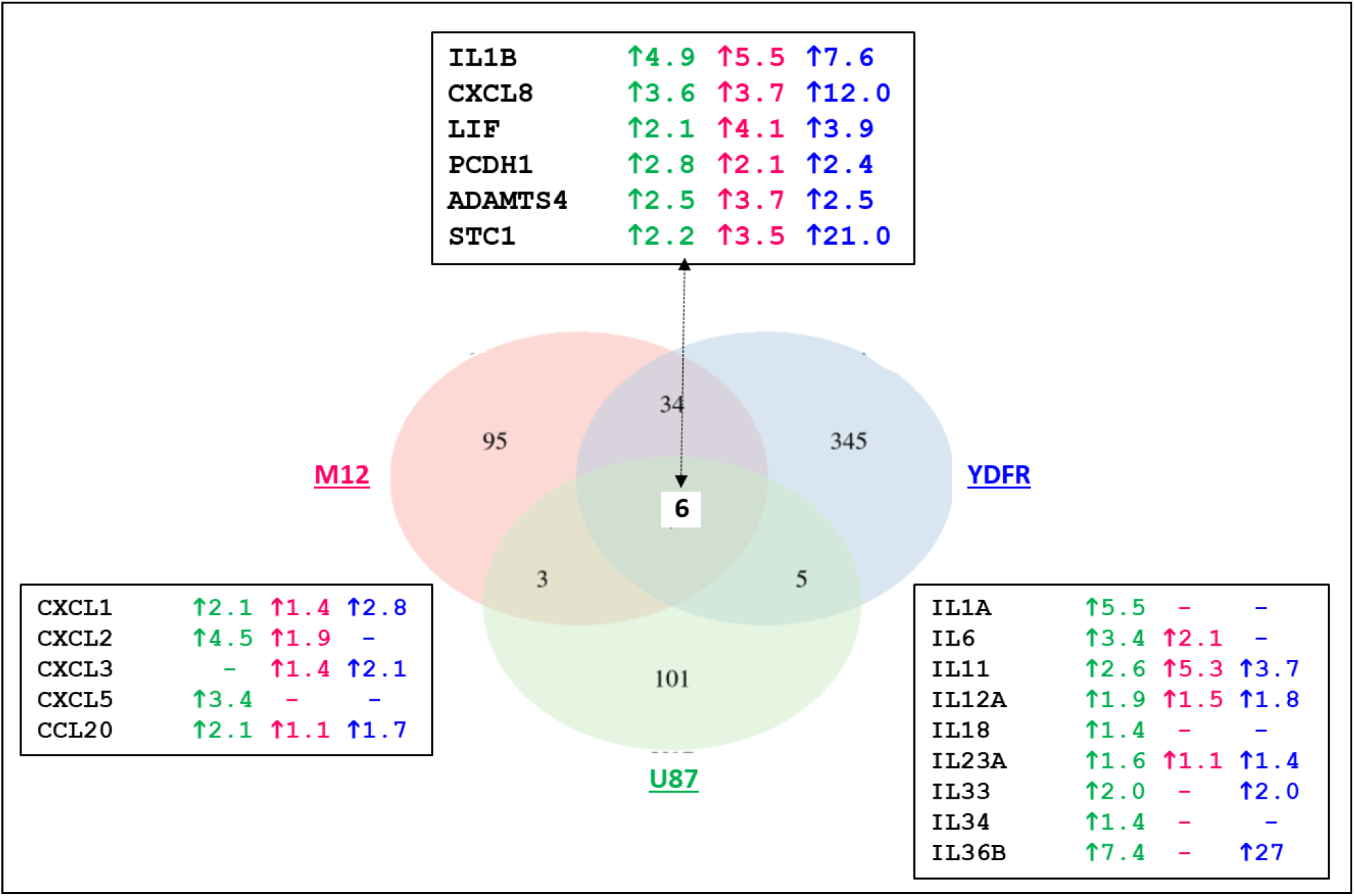
A comparison of the differentially expressed genes in the U87, M12 and YDFR cell lines that express the LY6S-iso1 protein. The upregulated genes in the cells expressing the LY6S-iso1 protein as compared to their respective control non-LY6S-iso1 expressing cells are represented as a Venn diagram, using the stringent criteria of p.adj<0.05, maximum counts>30 and a fold-change >2. The actual fold change for each cell line for the six genes that are upregulated in, and common to, all cell lines is shown in the top box (green, red and blue fonts representing the U87, M12 and YDFR cells, respectively). The fold changes for certain chemokines (CXCL and CCL) and interleukin (IL-) proteins are shown in the bottom left and bottom right boxes (dashes indicate no gene expression).

Accumulation of leukocytes (6.75E-17), (h) Cytokines and chemokine mediated signaling pathways (3.29E-16) and (i) Accumulation and recruitment of granulocytes (2.2E-14) and phagocytes (1.76E-14). These pathways reflected the alterations in gene expression patterns elicited by LY6S-iso1 expression, particularly of genes coding for cytokines, chemokines, chemokine receptors, interferon and immune and inflammatory related proteins (Fig. 10 and Table S2). Cytokines and chemokines pivotal for local and systemic responses to both immunological challenges and insults resulting from injury and infection that were upregulated, included IL6, IL36B, CXCL2, Granulocyte-Macrophage colony-stimulating factor (CSF2) and Granulocyte colony-stimulating factor (CSF3) (Fig. 10).

These results were confirmed at the protein level with cytokine/chemokine arrays, that showed increased secretion by the LY6S-iso1 expressing U87 cells of CXCL1, IL-1ra, CSF2, IL6 and MIP-1a (Fig. S11).

That LY6S-iso1 elicits expression of genes associated with an inflammatory phenotype was substantiated in the additional human cell lines, M12 and YDFR. In each of the three cell lines investigated (M12, YDFR and U87), LY6S-iso1 elicited increased expression (using the stringent criteria of p.adj <0.05, counts >30 and a fold-change>2) of IL1B, CXCL8, LIF, PCDH1, ADAMTS4 and STC1, all genes associated with an inflammatory cell phenotype (Fig. 11). Less stringent criteria showed that in at least two out of the three cell lines investigated LY6S-iso1 elicited increased expression of the chemokines CXCL1, CXCL2, CXCL3 and CCL20 and of the interleukins IL6, IL11, IL12A, IL23A, IL33 and IL36B (Fig. 11).

The following inflammation-associated genes (see Table S2) were also upregulated in all three LY6S-iso1-expressing cell lines-NRG1 (7.7-fold, 42-fold and 2.7-fold for M12, YDFR and U87 cells respectively), PTGS2 (1.6, 11 and 3.9), RCAN1 (1.5, 1.5 and 2.1) and SOCS1 (1.2, 1.9 and 2.8). Moreover, for all three [LY6S-iso1 expressing] cell lines the following canonical functional networks appeared in the top 14 pathways identified using the Ingenuity Pathway Analysis-IPA (Table S3a)-Dendritic Cell Maturation (2.236, 2.449 and 3.545, z-scores for M12, U87 and YDFR, respectively), IL-6 Signaling (2, 1.89 and 2.121), Acute Phase Response Signaling (2, 1.663 and 1.941), IL-8 Signaling (2.236, 1.342 and 1.291) and Neuroinflammation Signaling (1.633, 1.342 and 1.512). The IPA-derived Upstream Regulator Analysis provided further support for involvement of the LY6S-iso1 protein with an inflammatory cell phenotype. The list of Upstream Regulators (Table S3b) included the following top z-scoring Upstream Regulators for all three cell lines expressing LY6S-iso1 - TNF (number #1 in the list, with z-scores of 3.3, 5.0, 5.3 for M12, U87 and YDFR cells respectively), TPA-tetradecanoylphorbol acetate (#2, 3.6, 4.4 and 4.3), IL1B (#3, 2.4, 5.1 and 3.6), LPS-lipopolysaccharide (#4, 2.9, 5.3 and 2.8), IL17A (#5, 2.2, 4.7 and 4.0), NFkB (complex) (#7, 3.0, 4.4 and 2.9), ERK (#10, 3.1, 3.2 and 3.4), IKBKB (#12, 2.7, 4.1 and 2.6) IL1A (#13, 3.2, 3.7 and 2.7) and IL1 (#16, 3.2, 3.7 and 2.4). Additional Upstream Regulators included poly rI:rC-RNA (#20, 2.4, 3.8 and 2.9), TGFB1 (#23, 2.4, 2.2 and 4.1), TLR3 (#25, 2.5, 3.6 and 2.5) and TLR9 (#26, 2.4, 3.8 and 2.5).

## Discussion

The present study documents a previously un-annotated human gene now designated *LY6S*. Analyses both at the RNA and protein levels as well as TransMap and public domain RNA-Seq data provide evidence for the existence of the *LY6S* gene. At the RNA level, cloning and sequencing of the cDNA products generated by RT-PCR of human spleen samples showed mRNAs that code for three LY6S isoforms-LY6S-iso1 codes for a cell-surface protein, whereas LY6S-iso2 and LY6-iso3 code for secreted proteins. LY6S-iso1 has the ‘fingerprint’ features characteristic of the large LY6 protein family including (i) spacing of cysteine residues, (ii) exon/intron makeup in which exon 1 codes for the Signal Peptide, exon 2 codes for cysteines 1-5, and exon 3 codes for cysteines 6-10, (iii) the cysteine-cysteine (CC) doublet followed by (iv) the cysteine-asparagine (CN) doublet. What differentiates it from other LY6 family members is the consensus sixth cysteine residue replaced in LY6S-iso1 by a serine residue. We are unaware of any other member of the LY6 protein family that shows this deviation from the LY6 consensus. Because the unpaired, non-disulfide cysteine residue is now free to form disulfide bonds with unbonded cysteine residues in other proteins, the odd number (9) of cysteine residues in LY6S-iso1 likely affects LY6S-iso1 protein interactions. Yet 9 of the 10 consensus LY6S cysteine residues are retained in the LY6-iso1 protein, indicating that the four disulfide bridges formed between the LY6 consensus cysteine (C) residues C#1 and C#5, C#2 and C#3, C#7 and C#8 and C#9 and C#10 (*13*) should all be intact. Snake toxins that contain eight cysteine residues, associated with four disulfide bridges, form the characteristic three-finger structure (*2*) as do the LY6 proteins that have 5 disulfide bridges (*26*). Furthermore the amino-terminal LU/uPAR domain of uPAR lacks a cysteine pair (*27*), yet still forms the LU/uPAR domain. It is reasonable to assume, therefore, that the LY6S-iso1 protein also adopts a TFP structure.

Although the data indicate that human *LY6S* is homologous to the Ly6a subfamily of LY6 genes, we do not know whether *LY6S* is the ortholog of a single gene of the murine Ly6a subfamily, or whether all 8 murine genes and their protein products are an expansion of the solitary human LY6S gene. The phylogenetic analysis as well as our additional similarity analyses show that the eight murine Ly6 genes constitute a gene cluster, in which an ancestral gene likely underwent several duplication events, thereby forming the Ly6a subfamily cluster. Moreover, the TransMap algorithm of the UCSC genome browser maps *Ly6a* to the human *LY6S* gene (Fig. S1). These considerations lead us to conclude that this is a one to many orthologous relationship and *LY6S* is the human ortholog of the Ly6a subfamily genes.

Genes of the murine Ly6a subfamily play pivotal roles both in inflammation and in hematopoietic stem cells (HSCs). Murine *Ly6c* expression differentiates between two distinct types of monocytic/macrophage-like cells, and is markedly upregulated in inflammatory cells (for example (*28-32*)), and many of the Ly6a subfamily genes are related both to an inflammatory cell phenotype and to interferon related pathways (*21, 22*). In ADAR1 deficient mice in which extreme activation of interferon signaling pathways is observed, *Ly6a* and *Ly6c1* are upregulated 100- and 18-fold respectively (*20*), and *Ly6c2*, *Ly6i*, *Ly6g2* and *Ly6a2*, all members of the Ly6a subfamily, are also upregulated.

Just like the genes of the Ly6a subfamily, it appears from our analyses that *LY6S* is also involved in inflammatory processes. RNA-Seq analyses in three different human cell lines showed that expression of the LY6S-iso1 protein is associated with the expression of genes coding for chemokines, cytokines and of other proteins classically connected to an inflammatory cell phenotype. These findings lend further support, in addition to that provided by the phylogenetic and protein similarity analyses, to the notion that *LY6S* has a 1 to many orthologous relationship with the genes of the murine Ly6a subfamily gene cluster.

Like the genes of the Ly6a subfamily, the human *LY6S* gene is also an interferon stimulated gene (ISG). By inducing expression of hundreds of genes, interferons and the protein products of the induced ISGs are critical players in the restriction of viral infections. In this context murine *Ly6e*, the Ly6 gene immediately flanking upstream the Ly6a subfamily cluster (see Fig. 1, panel a) is a well-known ISG (*33*), that has recently been shown to differentially regulate infection with dependence on virus, acting on the one hand as a pan restriction factor of zoonotic coronaviruses (*34*), while enhancing infection of Influenza virus, West Nile virus or VSV, on the other (*35*) (*36*).

In line with our findings that *LY6S* expression is both interferon-inducible and associated with increased expression of genes related to inflammation and immune responses, we observed that LY6S-iso1 expression markedly inhibited viral replication, a phenomenon seen in each of the three different human cell lines investigated. This is likely indirectly mediated by LY6S-iso1 which, as noted above, leads to altered patterns of gene expression, that in turn affects virus replication. Thus, one of the functions of *LY6S* may be to elicit protection from viral infection, as already noted for *LY6E (34)*.

*LY6S* expression at the RNA level was highest in spleen whereas it was undetectable in bone marrow and peripheral blood leucocytes. At the protein level immunostaining analyses identified the LY6S-iso1 protein in about 5% of all spleen cells, yet it could not be detected in classical lymphoid-rich tissues such as thymus and tonsil. These findings suggest that the LY6S+ cells are likely not of a known classical hematopoietic or lymphoid cell lineage. In this respect, the expression pattern of *LY6S* in cells and tissues is different to that of murine *Ly6a* which, in contrast to *LY6S*, is expressed in peripheral blood leucocytes, on lymphoid precursor cells and hematopoietic stem cells in the mouse bone marrow (*37, 38*), and in many CD4+ T cells in the spleen.

In line with the notion that LY6S expression does not designate a classical cell of the human myeloid lineage, immunofluorescent analysis for expression of CD11b, a prototypical marker of human macrophages and possibly for many cells of human myeloid lineages, failed to show significant colocalization in the spleen with LY6S-expressing cells, just as classical T- and B-cell markers also failed to colocalize with LY6S expression. This should be viewed however in the following context-regarding the LY6S^+^ spleen cells about 10% of all CD45+ cells, a pan-leukocyte marker, are also LY6S^+^ (see Fig. 7, panels d), and a very few CD11b^+^ cells, amounting to less than 1% of these cells, are also LY6S^+^. Certain subsets of spleen macrophages, such as red pulp, marginal zone, and marginal metalophillic macrophages do not express high levels of CD11b if at all, and because of this the LY6S positive cells may belong to such a subset.

In an attempt to learn more about the LY6S-iso1^+^ spleen cells, we queried publicly available human spleen single-cell RNA-Seq data but because of their limited gene coverage and low sensitivity, LY6S-iso1^+^ cells could not be identified in the available single cell RNA-Seq data sets. Sorting of human spleen cells with the anti-LY6S-iso mAbs that we have generated into LY6S-iso1+ve cells and LY6S-iso1 negative cells, followed by RNA-Seq analyses might assist in the future identification of the [LY6S-iso1^+^] cell type.

In summary, we have identified the new interferon-inducible human *LY6S* gene that is likely the long-sought human ortholog of genes belonging to the *Ly6a* subfamily. *LY6S* is expressed in a discrete subset of human spleen cells of a non-classical lineage, and its expression is associated with an inflammatory cell phenotype and with the restriction of viral replication. Of particular therapeutic clinical interest are the recent findings showing that Ly6e and Ly6a serve as receptors for viral entry into the cell. Ly6e is a receptor for HIV (*39*) and Ly6a, expressed on the surface of murine brain endothelial cells, is the cell receptor for a recombineered Adeno Associated Virus (AAV-PHP.B), via which AAV-PHP.B can penetrate the Blood Brain Barrier and transduce gene cargo into the brain (*40-44*). Our findings indicate that human LY6S is the ortholog to the genes of the murine *Ly6a* subfamily, and as we have observed LY6S expression in human brain tissue, the discovery of *LY6S* as reported here may open up new possibilities for introducing therapeutic genes into the human brain. Furthermore, the LY6S proteins may serve as receptors for additional pathogenic viruses, thus leading to novel anti-viral therapeutic strategies.

## Methods

### Reagents and antibodies

All chemicals and reagents were obtained from Sigma (St. Louis, MO), unless otherwise specified. Secondary antibodies used in cell counter-staining or in immunohistochemical development were obtained from Jackson ImmunoResearch Laboratories (Bar Harbor, ME).

### Immunization of mice and generation of hybridomas

Mice were immunized with 5 consecutive intradermal immunizations spaced at 21-day intervals, with peptide E, derived from the amino acid sequence of LY6S-iso1 (LY6S-iso1-pepE, see Fig. 6a), covalently conjugated to Keyhole Limpet Hemocyanin (KLH-pepE). The mice were purchased from Harlan Laboratories Limited (Israel) and were housed and maintained in laminar flow cabinets under specific pathogen-free conditions in the animal quarters of Tel-Aviv University. All work was performed in accordance with current regulations and standards of the Tel-Aviv University Institutional Animal Care and Use Committee. The initial immunization was done with Freund’s adjuvant, and the following 4 KLH-pepE boost immunizations together with incomplete Freund’s adjuvant. Hybridomas were prepared by fusion of nonsecreting myeloma cells with immune splenocytes and screened by ELISA assay (see below).

### ELISA for determining binding of anti-LY6S-pepE monoclonal antibodies to LY6-iso1 protein

Elisa Immunoassay plates (CoStar) were coated with recombinant hFc-LY6-iso1 protein, produced and secreted by HEK 293 cells stably transfected with and secreting hFc-LY6S-iso1 protein (see below here-‘Expression of hFc-LY6S fusion protein’). Spent culture media from the initial hybridomas was then applied to the wells. Following incubation, samples were removed and the wells were washed with PBS/Tween. Detection of bound antibodies was done with horseradish peroxidase (HRP)-conjugated anti-mouse antibody.

### Expression of hFc-LY6S fusion protein

The hFc-LY6S fusion protein was generated by inserting DNA coding for this protein into the pcDNA3.1 expression vector using standard molecular biology techniques. The make-up of this insert, the amino acid sequence of the resultant fusion protein and the DNA sequence of the insert are shown in Fig. S12.

### Sequence determination of SCA5E12 anti-LY6-iso1 monoclonal antibodies

RNA was isolated from the SCA5E12 hybridoma (anti-LY6-iso1 mAb) with TRIzol® Reagent l, according to a technical manual for the reagent (Ambion Inc., Foster City, CA). The sequence of the SCA5E12 antibody was determined as follows: cDNA was generated by reverse transcription of total RNA with the use of universal or isotype-specific anti-sense primers, according to the technical manual for PrimeScriptTM First Strand cDNA Synthesis Kit (Takara Bio Inc., Mountain View, CA). Amplification of VH and VL antibody fragments was carried out according to the standard operating procedure (GenScript, NJ, USA) which involves rapid amplification of cDNA ends, followed by separate cloning into a standard cloning vector. Clones with inserts of the correct sizes were sequenced by colony PCR and at least five colonies with such inserts were sequenced for each fragment, with the consensus sequence derived by alignment of the different clones.

### Immunohistochemistry

Chromogenic immunolabeling for SCA5E12 was performed on formalin-fixed, paraffin embedded human tissue sections. Briefly, following dewaxing and rehydration, slides were immersed in 1% Tween-20, then heat-induced antigen retrieval was performed in a pre-heated steamer using Antigen Unmasking Buffer (catalog# H-3300, Vector Labs) for 25 minutes. Slides were rinsed in PBST and endogenous peroxidase and phosphatase was blocked (catalog# S2003, Dako) and sections were then incubated with primary SCA5E12 mouse monoclonal antibody (1:50 dilution) for 45 min at room temperature. (For staining in the presence of competing peptides, primary antibody was diluted 1:10 in the peptide solution directly.) The primary antibodies were detected by 30 minute incubation with HRP-labeled anti-mouse secondary antibody (catalog# PV6114, Leica Microsystems) followed by detection with 3,3′-Diaminobenzidine (catalog# D4293, Sigma-Aldrich,), counterstaining with Mayer’s hematoxylin, dehydration and mounting.

### Immunofluorescence

Dual OPAL immunofluorescent labeling with SCA5E12 and anti-CD45 antibodies (catalog# 145M-94, Cell Marque/Sigma Aldrich), was performed on formalin-fixed, paraffin embedded tissue sections following manufacturer’s instructions (NEL810001KT Akoya Biosciences, Menlo Park, CA). Briefly, following standard dewaxing and rehydration, slides were immersed in Antigen Unmasking Buffer (H-3300 Vector Laboratories, Burlingame, CA) in a plastic chamber and retrieved in a microwave, which is also the subsequent antibody stripping step for sequential multiplex staining. Endogenous peroxidase and phosphatase was blocked (catalog# S2003, Dako) and sections were then incubated sequentially with each primary antibody (1:80 dilution for SCA5E12 and specific dilutions for the other primaries as listed below) for 45 minutes at room temperature with antibody stripping step performed in between. Slides were incubated with differently labeled anti-Mouse IgG (PV6114 Leica, Wetzlar, Germany) or anti-rabbit IgG (PV6119 Leica) secondary antibodies, as appropriate, for 30 minutes. Fluorescent dyes OPAL 520 (for each of the antibodies listed below) and OPAL 690 (for SCA5E12) were diluted in OPAL amplification buffer and slides were stained for 10 minutes. Slides were counterstained with DAPI working solution for 10 minutes, washed, and mounted with ProLong Gold.

Information on the specific primary antibodies used for dual IF with the SCA5E12 antibody: Anti-CD4: ab133616 / 1:200/Rabbit/Abcam; Anti-CD8: m0814/1:100/Mouse/Dako; Anti-CD11b: ab133357/ 1:8000/Rabbit/Abcam; Anti-CD11c: ab52632/ 1:100 /Rabbit/Abcam; Anti-CD19 : catalog# 119M-14,/1:50/Mouse/Cell Marque/Sigma Aldrich; Anti-CD20: 0755/1:100/Mouse/Dako; Anti-CD31: RB-10333-P/1:50/Rabbit/Thermo Fisher Scientific; Anti-CD34: catalog# 134M-14,/1:75/Mouse/Cell Marque/Sigma Aldrich; Anti-CD45: catalog# 145M-94,/1:100/Mouse/Cell Marque/Sigma Aldrich; Anti-CD68: m0814/1:250/Mouse/Dako; Anti-CD117 (c-Kit): A450229-2/ 1:400/Rabbit/Dako; Pan-cytokeratin AE1/AE3: ab961/1:10/Mouse/Abcam; Anti-FoxP3: 12653S/1:50/Rabbit/Cell Signaling Technologies.

Samples used for immunohistochemical stainings were procured from the Biomax tissue bank, with all required ethical approvals therein, as stated in http://biomax.us – “All tissue is collected under the highest ethical standards with the donor being informed completely and with their consent. We make sure we follow standard medical care and protect the donors’ privacy. All human tissues are collected under HIPPA approved protocols”.

### Generation of pQCXIP plasmids coding for LY6S proteins and generation of stable cell infectants expressing the LY6S proteins

Total RNA was extracted from human melanoma M12-CB2 cells that were treated with gamma interferon for 48 hours using the EZ-RNA Total RNA Isolation Kit (Biological Industries, Kibbutz Beit-Haemek, Israel), followed by cDNA synthesis with qScript cDNA Synthesis Kit (Quantabio). The over-expression construct of human LY6 was created by PCR amplification of cDNA by Phusion^®^ High-Fidelity DNA polymerase (Thermo Fisher Scientific), using the following primers LY-6S-Pac1-S-5’-GGAACGTTAATTAACGCTGAAGTTTGTCTGTGCACTA-3’, LY6S-EcoR1-AS-5’-AGAGAGGAATTCCCAAGCAGCTGGTGACGCACAG-3’. The generated fragment was digested with Pac1 and Ecor1 and ligated into the corresponding sites of pQCXIP vector (Clontech Laboratories, Inc., Mountain View, CA, USA). PCR products of LY6S were sequenced. Cells were infected to stably express LY6S-pQCXIP retroviral vector (Clontech Laboratories, Inc., USA) as described previously (*45*).

### Vesicular Stomatitis Virus (VSV) infection

M12, YDFR or U87 cells were infected with VSV-New Jersey (a gift of Prof. Moshe Kotler, Hebrew University) at multiplicities of infection of 0.5,0.15 or 0.05 for 24 or 48 hours (as indicated in Fig. legend). Titer of inoculum or of spent medium of infected cells was measured by plaque assay on Vero cells: sequential 10-fold dilutions in DMEM, 125,000 cells/well per well of 12 well plate. Overlay was with 0.6% methylcellulose. Detection was by crystal violet staining.

### Flow cytometry

Antigen expression was determined using Flow cytometer S100EXi (Stratedigm, San Jose, CA) with CellCapTure software and FlowJo v10. Dead cells were gated out from the analysis.

### Phylogenetic Tree Analysis

Sequences were aligned by using the Muscle Multiple Sequence alignment software (PMID:30976793) available through an online server at the EBI (https://www.ebi.ac.uk/Tools/msa/muscle/). AliView (PMID:25095880) was used to view and edit the initial alignment of 303 columns. The alignment was edited to remove indel regions of ambiguous alignment and columns with <20% gaps were retained, providing a final alignment of 113 amino acids.

ProtTest-3.4.2 (*46*) was used to determine the best fit evolutionary model for this maximum likelihood analysis. The RAxML-NG (PMID: 31070718) blackbox server (https://raxml-ng.vital-it.ch/#/) was used to perform a maximum likelihood analysis (JTT+G4m substitution model, stationary base frequencies taken from the model, with the proportion of invariant sites box selected, and four gamma rate categories to model the among-site rate heterogeneity), with automatic bootstopping of the bootstrapping procedure, as implemented in RAxML. Trees were visualized and edited with FigTree v1.4.3 software (http://tree.bio.ed.ac.uk/software/figtree/).

### mRNA-seq

Libraries were prepared by the Crown Institute for Genomics (G-INCPM, Weizmann Institute of Science) using an in-house poly A based RNA seq protocol (INCPM-mRNA-seq). Briefly, the polyA fraction (mRNA) was purified from 500 ng of total input RNA followed by fragmentation and the generation of double-stranded cDNA. After Agencourt Ampure XP beads cleanup (Beckman Coulter), end repair, A base addition, adapter ligation and PCR amplification steps were performed. Libraries were quantified by Qubit (Thermo fisher scientific) and TapeStation (Agilent). Sequencing was done on a Nextseq 75 cycles high output kit, allocating 20M reads per sample (Illumina; single read sequencing). The RNA-Seq data, GEO accession GSE188924, has been uploaded to the GEO database. This SuperSeries record provides access to all of the data relating to both the control human cells (U87 glioblastoma, M12 melanoma and YDFR melanoma cells) and their counterparts expressing the LY6S-iso1 protein. To review it please go to: https://www.ncbi.nlm.nih.gov/geo/query/acc.cgi?acc=GSE188924

The Paired End (PE) True Stranded RNA-Seq data, GEO accession GSE159456 has also been uploaded to the GEO database at: https://www.ncbi.nlm.nih.gov/geo/query/acc.cgi?acc=GSE159456

### Data Analysis for RNA-Sequencing

Poly-A/T stretches and Illumina adapters were trimmed from the reads by using cutadapt; reads shorter than 30bp were discarded. Reads for each sample were aligned independently to human reference genome GRCh38 with STAR (*47*), and supplied with gene annotations downloaded from Ensembl (and with the EndToEnd option). Counting proceeded over genes annotated in Ensembl release 92 using the htseq-count (with stranded= “reverse” option for the PE Paired End) (*48*). Only uniquely mapped reads were used to determine the number of reads falling into each gene (intersection-strict mode). Differential analysis was performed with the DESeq2 package (*49*), the betaPrior, cooksCutoff and independentFiltering parameters set to False. Raw P values were adjusted for multiple testing by using the Benjamini and Hochberg procedure. Differentially expressed genes were determined by a p-adj of < 0.05 and absolute fold changes > 2 and max raw counts > 30. Functional analysis of DE genes was performed with IPA (QIAGEN Inc., https://www.qiagenbioinformatics.com/products/ingenuity-pathway-analysis), together with Martin, M. (2011, “Cutadapt removes adapter sequences from high-throughput sequencing reads” *EMBnet. Journal*, *17*(1), pp–10doi:<u>10.14806/ej.17.1.200</u>) and methods detailed in references (*47-50*).

## Author contributions

Discovery of LY6S: DHW, MS, MC

Immunofluorescent and immunohistochemical studies: AM, SR, MC

Analyses of RNA-Seq data sets: NS, AD, ASP, DHW

Anti-LY6S-iso1 monoclonal antibodies: DHW, OSA, RZ, NIS

Cell transfectants expressing LY6S proteins: TM, OSA, SL, BN, DB, AS, RZ, DHW

Viral resistance of LY6S-iso1 expressing cells: ML, ME

Phylogenetic analyses and gene annotation: EB, BB, DHW

Writing-original draft: DHW, EB, BB, NS, AD

Writing-review and editing: DHW, EB, ME, IPW

Funding acquisition: IPW, DHW

Project administration: DHW, IPW

Supervision: DHW, IPW

**Competing interests:** None to declare.

---

**Fig. S1.**
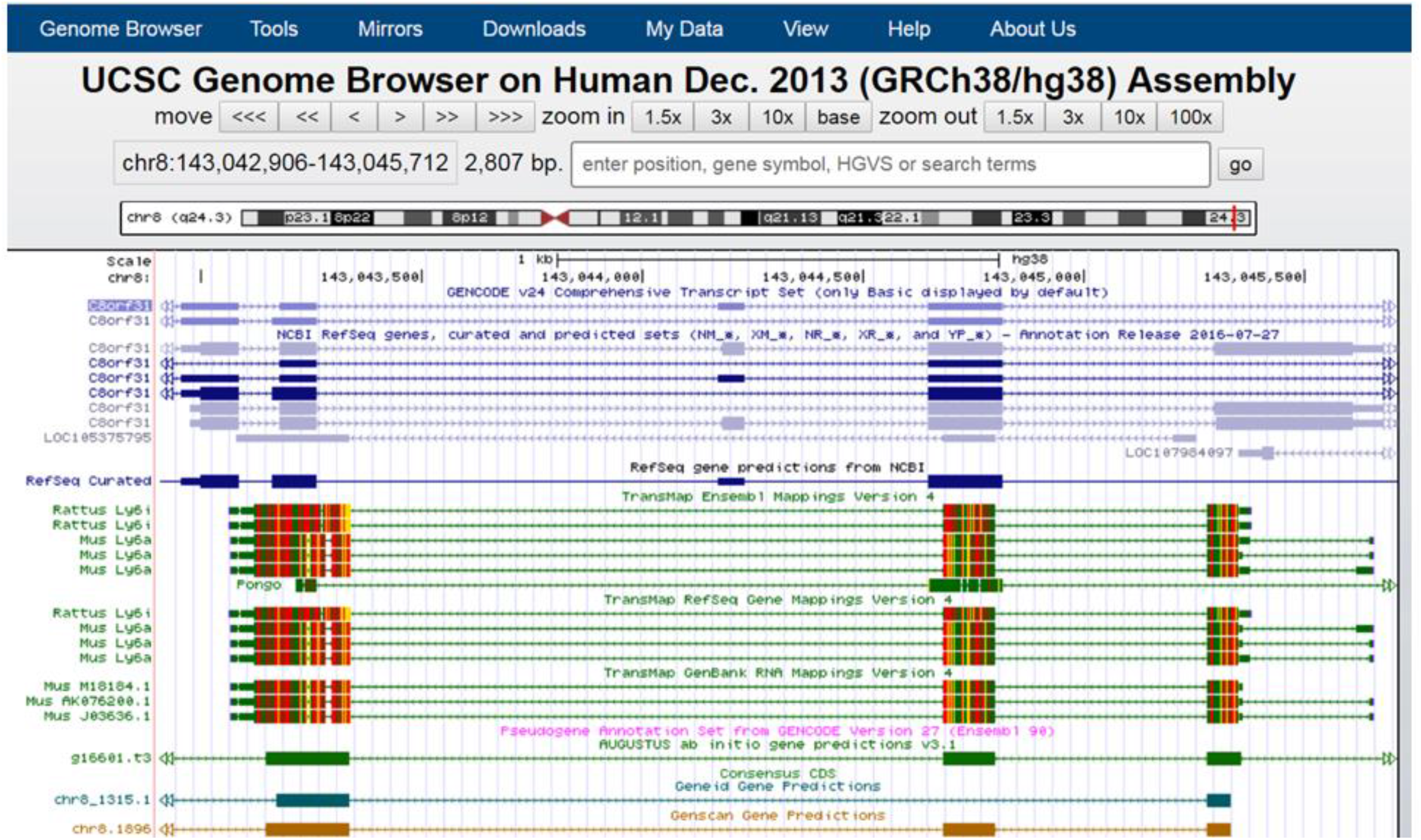
TransMap alignments position the murine Ly6a gene at the location of the predicted chr 8.1896 Genscan gene-human LY6S. TransMap, the cross-species alignment algorithm which maps genes and related annotations from one species to another by using synteny-filtered pairwise genome alignments to determine the most likely orthologs, maps mouse *Ly6a/Sca1* at the same position as *LY6S* shown here as the GenScan predicted human gene chr 8.1896. TransMap Ensembl Mappings Version 4, TransMap RefSeq Gene Mappings Version 4 and TransMap GenBank RNA Mappings Version 4 all map murine *Ly6a* (ENSMUST00000187994.6) to *LY6S*.

**Fig. S2.**
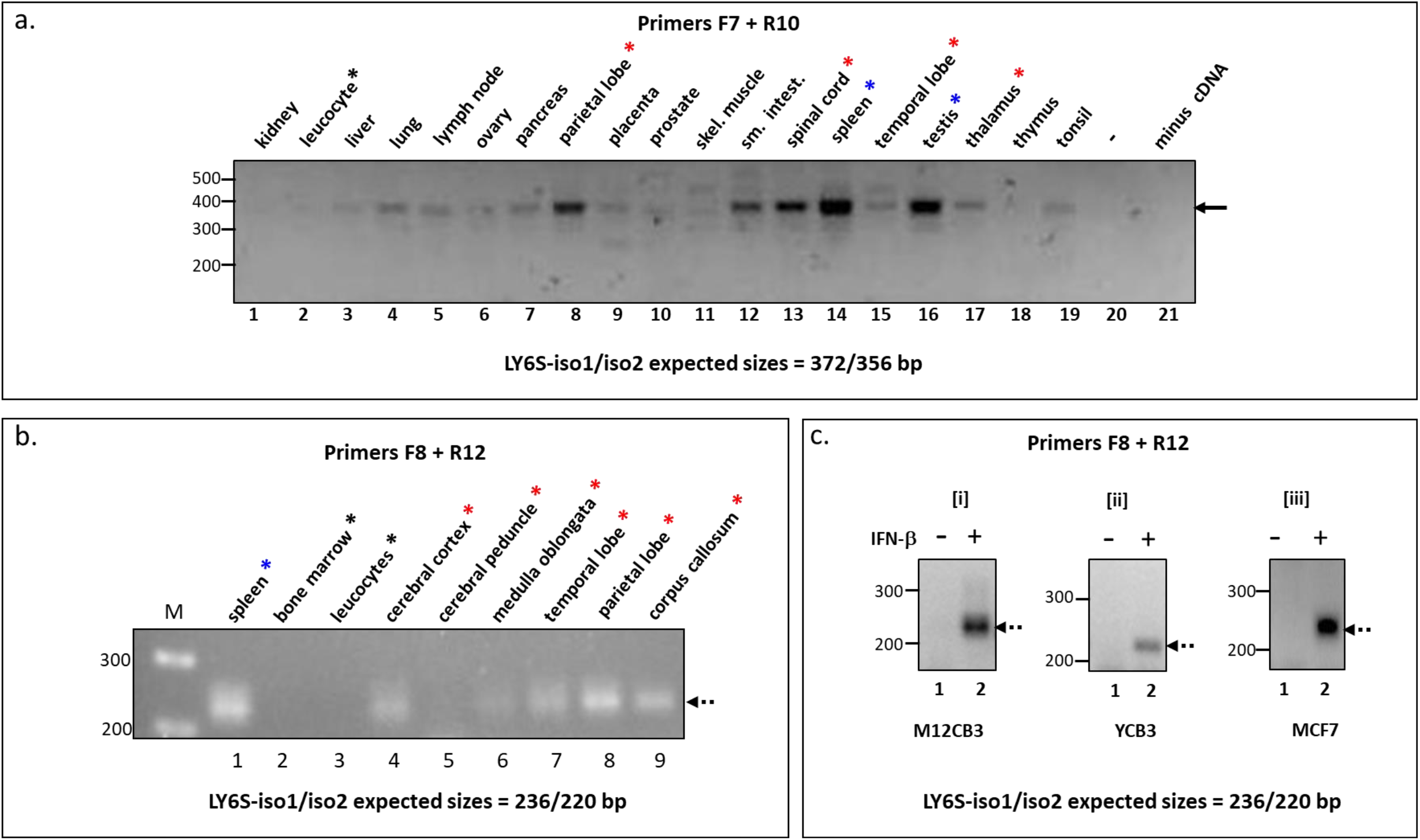
Expression of LY6S mRNA in various regions of the human brain and other tissues and its induction in cells by interferon. **(Panel a)** RT-PCR analysis of cDNAs derived from human tissues (40 cycles), done with LY6S primers F7 and R10 (see Table S1). The arrow to the right indicates the approximate position for the expected sizes of the LY6S RT-PCR products with these primers (372/356 bp for iso1 and iso2, respectively), and DNA marker sizes are shown at the left. Tissues from the central nervous system, are indicated by the red asterisks, spleen and testis by the blue asterisks and the negative leucocyte sample by the black asterisk. **(Panel b)** RT-PCR of additional regions of the brain done with LY6S primers F8 and R12. The arrow at the right indicates the approximate position for the expected sizes of the LY6S RT-PCR products with these primers (236/220 bp for iso1 and iso2, respectively). **(Panel c)** RT-PCR of cDNAs prepared from the melanoma cells M2CB3 and YCB3 (panels [i] and [ii] respectively) and from MCF7 cells (panel [iii]) done with LY6S primers F8 and R12 (see Table S1). The cells were either treated with interferon beta (lanes 2) or not treated (lanes 1), as indicated.

**Fig. S3.**
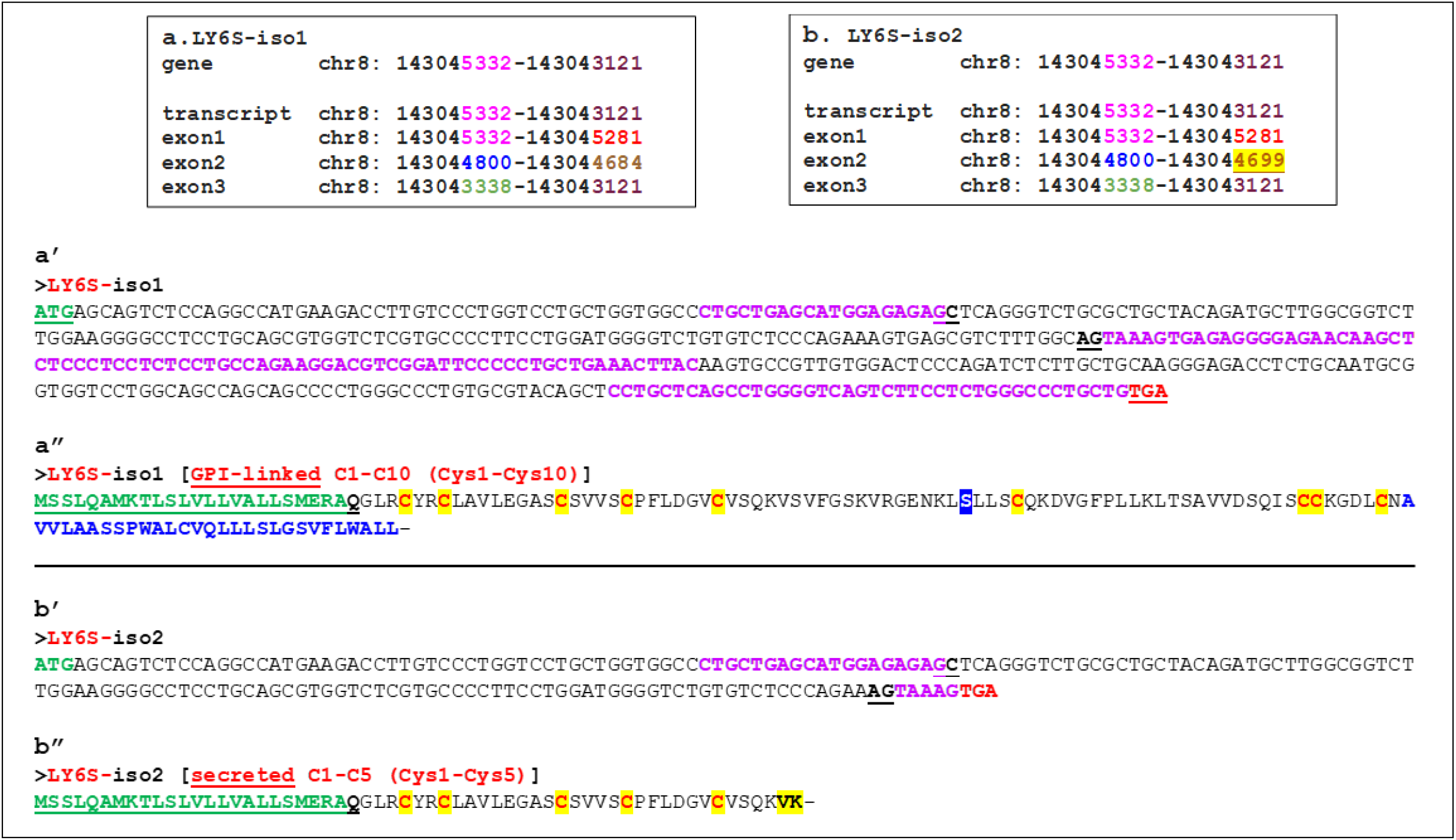
LY6S-iso1 and LY6S-iso2 genomic coordinates, nucleotide sequences, splice sites and amino acid sequences and nucleotide sequences which are unique for LY6S-iso1 and LY6S-iso2. **Panels a and b-** The genomic coordinates (GRCh38/hg38 Assembly) for LY6S-iso1 and LY6-iso2 mRNAs are shown. The *LY6S* gene is transcribed from the complementary strand so the directional coordinates go from high nucleotide numbers to lower nucleotide numbers. **Panels a’ and b’**-The nucleotide sequences of LY6-iso1 and LY6-iso2, respectively. Purple fonts are specific for *LY6S*, whereas regular black font sequences are common to the *LY6S* gene and the known *C8orf31* gene (see Fig. 1b, 1c and Fig. 3a and 3b). When searching for expression of the *LY6S* transcripts in RNA-Seq reads, the nucleotide strings shown in purple fonts must appear in the RNA-Seq sequences, because only these sequences are *LY6S* specific-the remaining sequences are common (complementary) to *C8orf31* transcripts. Splice sites are indicated by **<u>GC</u>** and **<u>AG</u>**. The initation and termination codons are in green and red underlined fonts, respectively. **Panels a” and b”**-Amino acid sequences of the LY6S-iso1 and LY6-iso2 proteins. The signal peptide (SP) in both LY6S-iso1 and LY6-iso2 is indicated in green, bold, underlined fonts, and the first amino acid of the mature, cleaved protein, as predicted by the Phobius SP predictor is the Q (glutamine) residue in black, bold font. Consensus cysteine (C) residues of LY6-like proteins present in LY6S-iso1 and LY6-iso2 are indicated in bold red fonts against a yellow background. In LY6S-iso1 the atypical S (serine) residue that replaces the LY6 consensus sixth cysteine is shown by a bold white font against a blue background. The C-terminal hydrophobic tail in LY6S-iso1 protein (panel a”) that initiates with the sequence AAVL…. (in blue bold fonts) provides the signal for addition of the GPI-anchor, as predicted with high probability and a 100% specificity by the PredGPI algorithm. The omega-site position to which the GPI-anchor attaches is the asparagine (N) residue immediately upstream to this C-terminal sequence. The C-terminal dipeptide sequence VK (valine-lysine) present only in LY6-iso2 and followed by a termination codon (panel-b”) is in bold black fonts against a yellow background.

**Fig. S4a.**
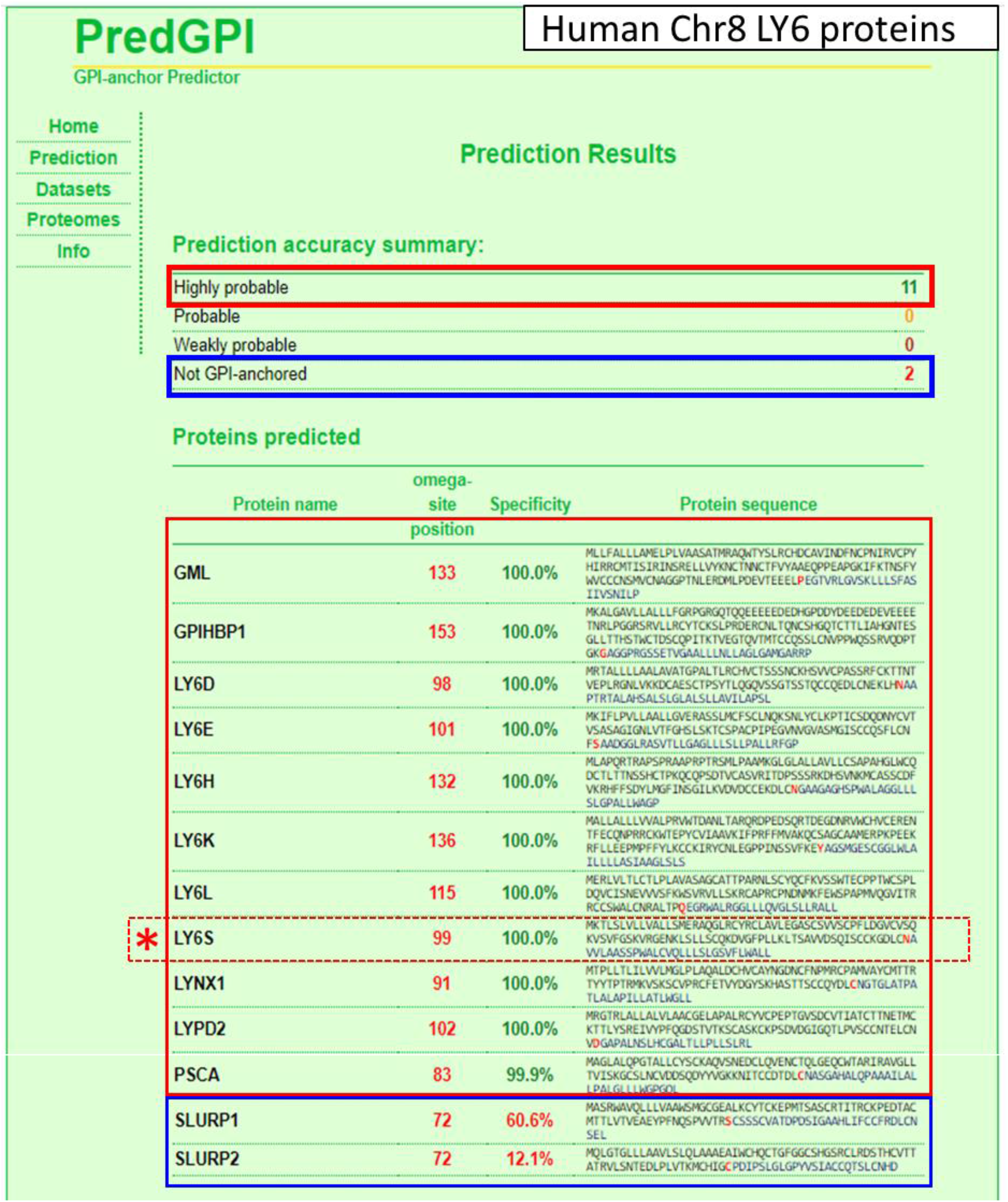
Prediction of GPI-linkage of human chromosome 8 LY6 *proteins*.

**Fig. S4b.**
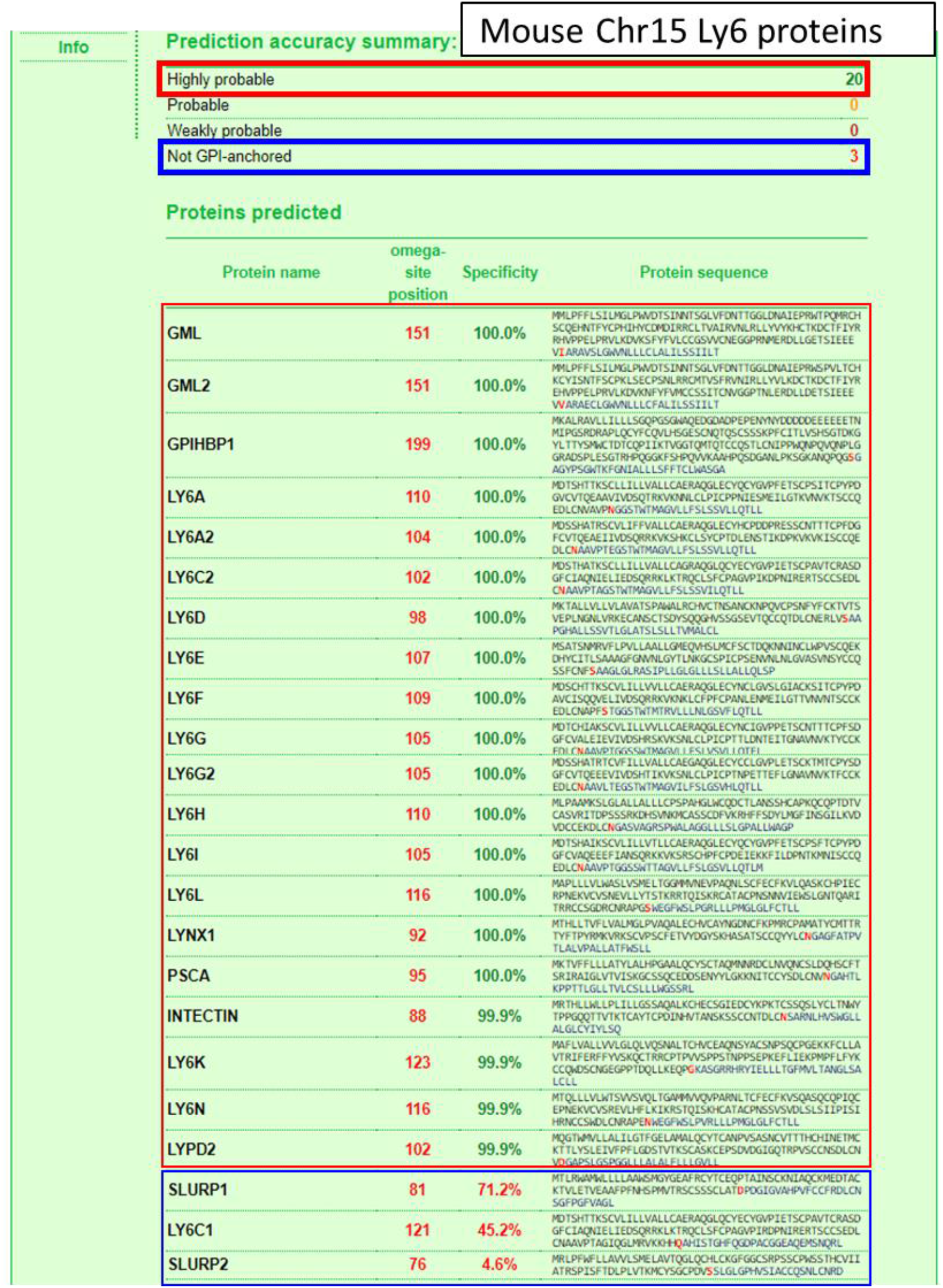
Prediction of GPI-linkage of mouse chromosome 15 LY6 proteins.

**Fig. S5.**
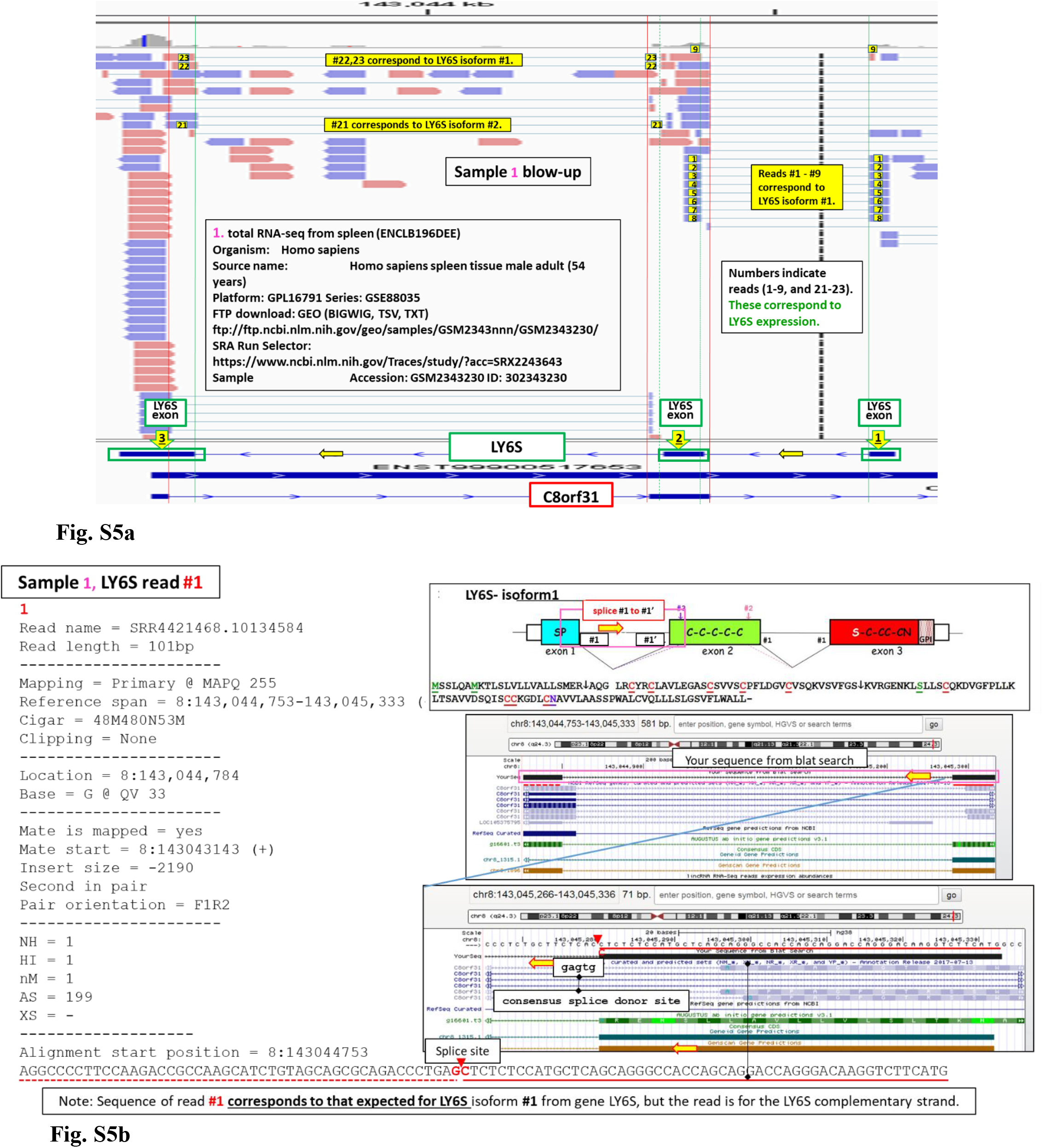
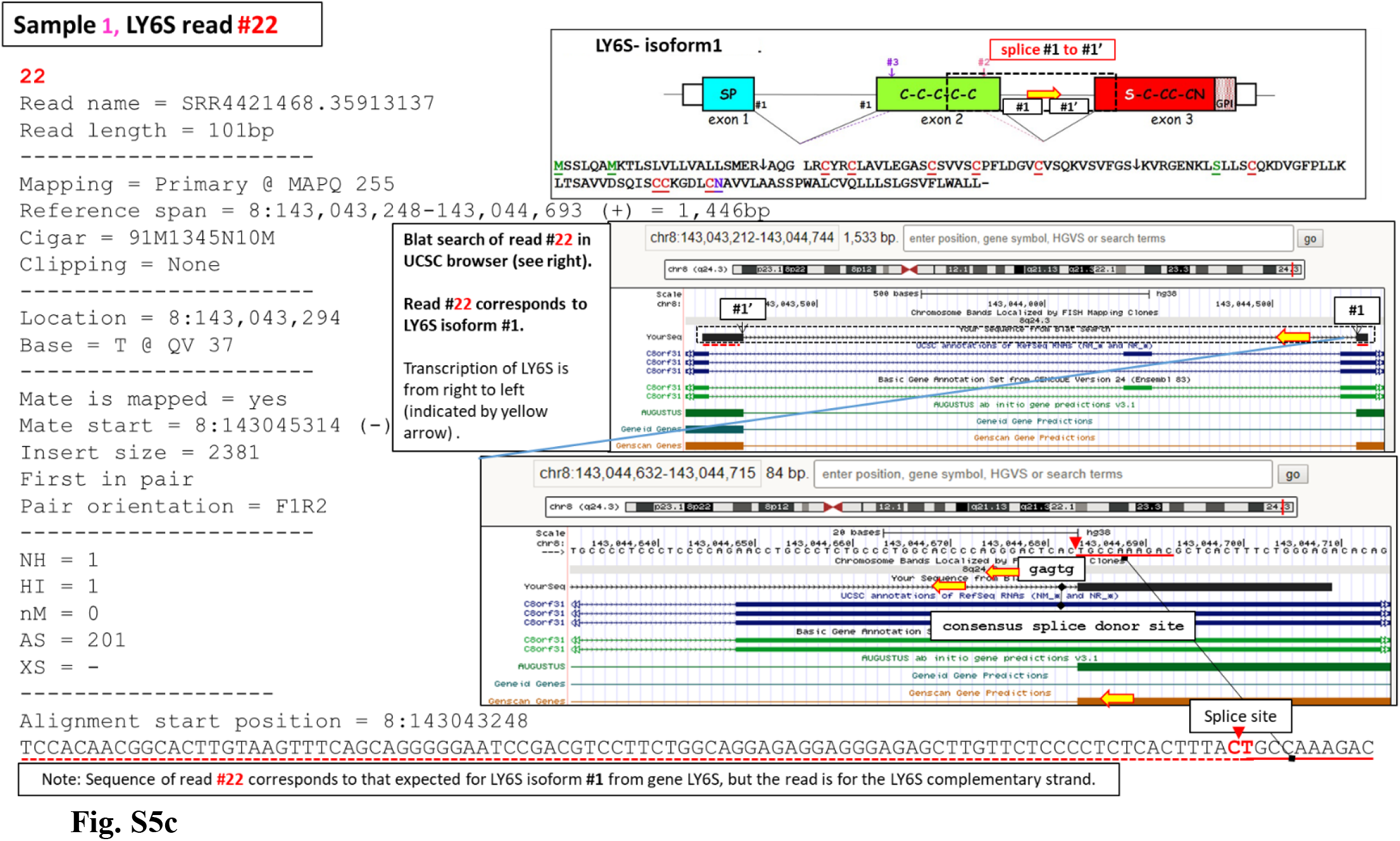
Analysis of the human spleen total RNA-Seq shows RNA reads that precisely correspond to transcripts derived from the LY6S gene. **(a)** The genomic coordinates of the human *LY6S* gene were introduced into our local gene database and the RNA-Seq data from human male spleen (Geo data set ENCLB196DEE) were queried against this modified database. The reads mapping to the selected region of the *C8orf31* and (newly added) *LY6S* genes were analyzed, and transcripts deriving from *LY6S* are indicated (numbers in black bold fonts against a yellow background). The *LY6S* direction of transcription is indicated by the horizontal yellow arrows; the *LY6S* exons and *LY6S* splice junctions are shown by the downward facing yellow arrows and vertical green lines, respectively. The vertical red lines show the *C4orf48* splice sites. (a) **and (c)-** Detailed analysis of representative reads deriving from LY6S-iso1 mRNA.

**Fig. S6.**
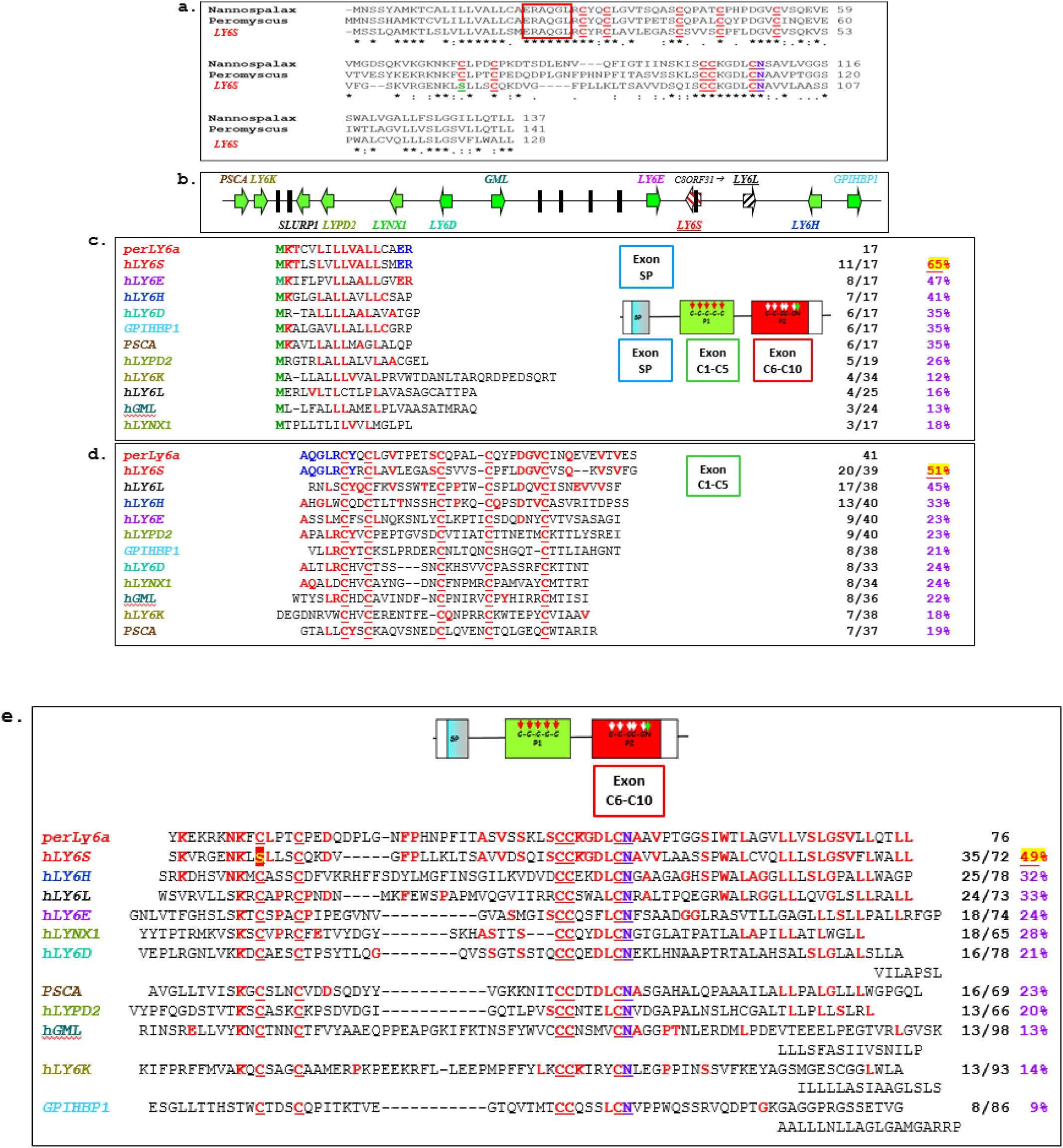
Sequence homology analysis of the Ly6a-2/Ly6e-1 (Ly6a) protein from the North American deer mouse (Peromyscus maniculatus bairdii) with human LY6S-iso1 and with all other LY6 proteins from the human chr8 LY6 locus. **Panel a**-The amino acid sequence of human LY6S-iso1 was compared with that of the Ly6a (synonymous with Ly6a-2/Ly6e-1) proteins from the North American deer mouse (Peromyscus maniculatus bairdii) and from Nannospalax galili. Identical amino acids in all three proteins are indicated by asterisks, similar amino acids by a colon, and the consensus ERAQGL peptide sequence forming the signature ‘fingerprint’ of all Ly6a subfamily cluster proteins is boxed by the red rectangle. **Panel b-** The locations of all human *LY6* genes (green arrows, transcriptional direction as indicated) on chromosome 8. LY6S is indicated by the red slashed arrow. **Panels c, d and e-** Homology comparison of Peromyscus maniculatus bairdii deer mouse Ly6a (designated perLY6a) protein with LY6S-iso1 and with all other LY6 proteins derived from the human chr8 *LY6* genes. The ‘ERAQGL’ sequence conserved in the C-terminal part of the signal peptides of the Ly6a subfamily cluster of genes cluster proteins is indicated in bold blue fonts; identical amino acids appearing in both perLY6-Sca-1 and hLY6S are in red bold fonts in both sequences. Amino acid residues in other chr8 LY6 proteins that are identical to perLy6a residues are in bold red fonts. The number of amino acid identities as well as the percent identity with the reference PerLy6a sequence is indicated at the right. The proteins are divided into sequences from Exon SP (exon 1), Exon C1-C5 (exon 2, coding for LY6 consensus cysteine residues 1-5) and Exon C6-C10 (exon 2, coding for LY6 consensus cysteine residues 1-5, panels c, d and e, respectively).

**Fig. S7.**
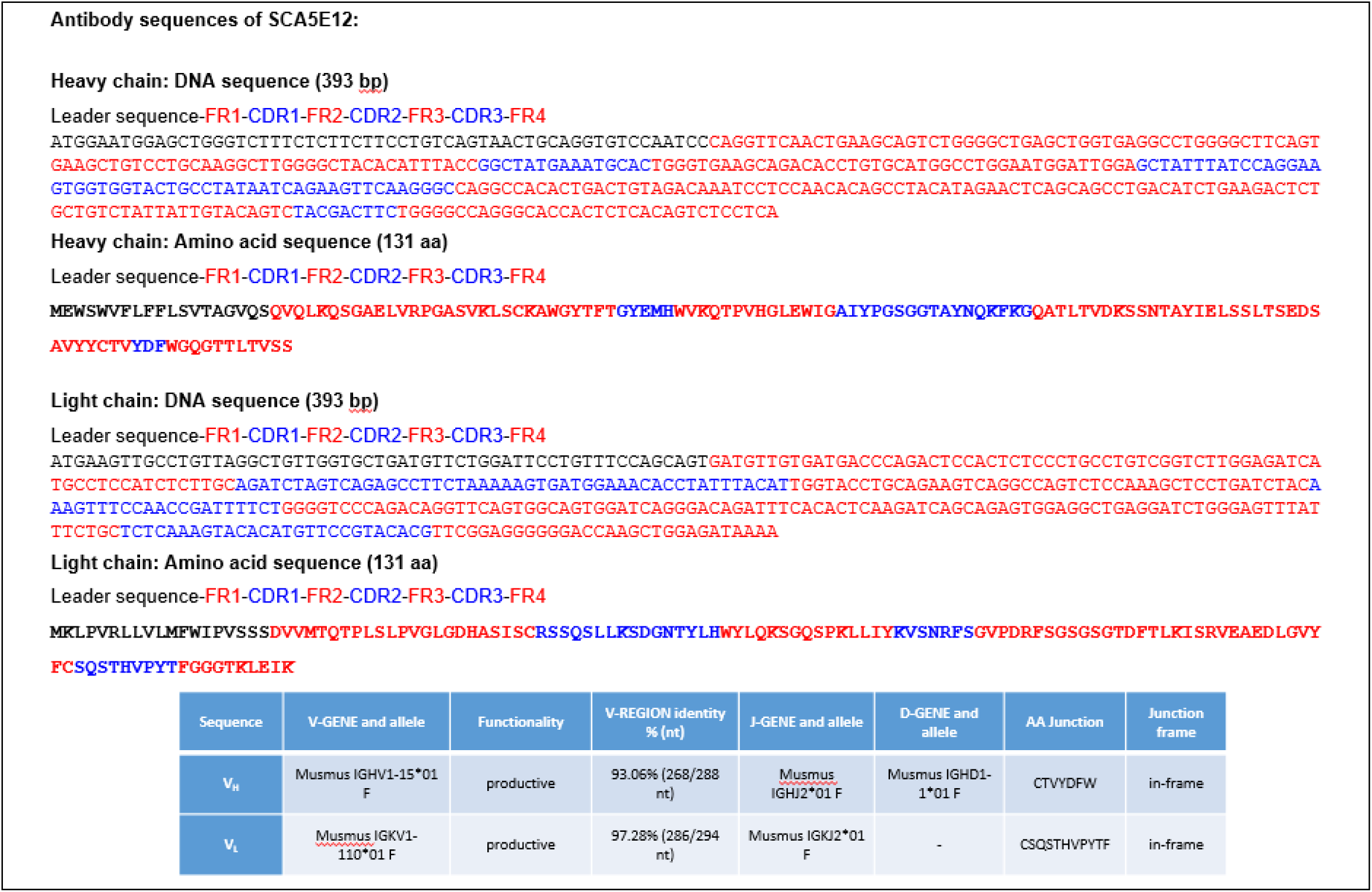
Amino acid and nucleotide sequences and genomic derivations of the variable domains of anti-LY6S-iso1 mAb-5E12. The sequences of the anti-LY6S-iso1 mAb-5E12 mAb were determined as described in Materials and Methods, and its genomic derivation is as shown at the top of the Fig. s.

**Fig. S8.**
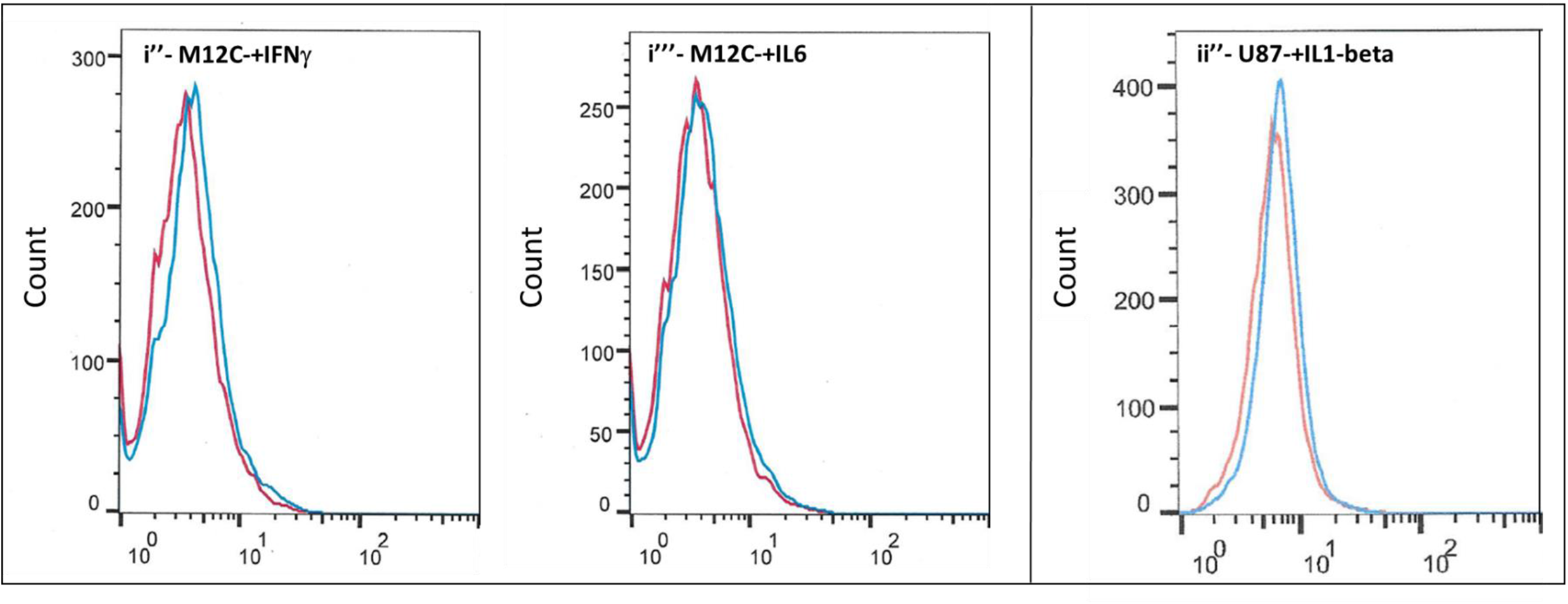
Detection of LY6S-iso1 protein with anti-LY6S-iso1 mAb 5E12 on the cell surface of cytokine treated melanoma and glioblastoma cells. Human melanoma (M12C, panels i” and i’”) or human glioblastoma (U87, panels ii”) cells were grown with interferon-beta (panel i”), IL6 (panel i”’) or IL1-beta (panel ii”) as indicated and assessed by flow cytometry with secondary antibody alone (red lines) or with mAb-5E12 followed by the fluorescently labelled anti-mouse secondary antibodies (blue lines).

**Fig. S9.**
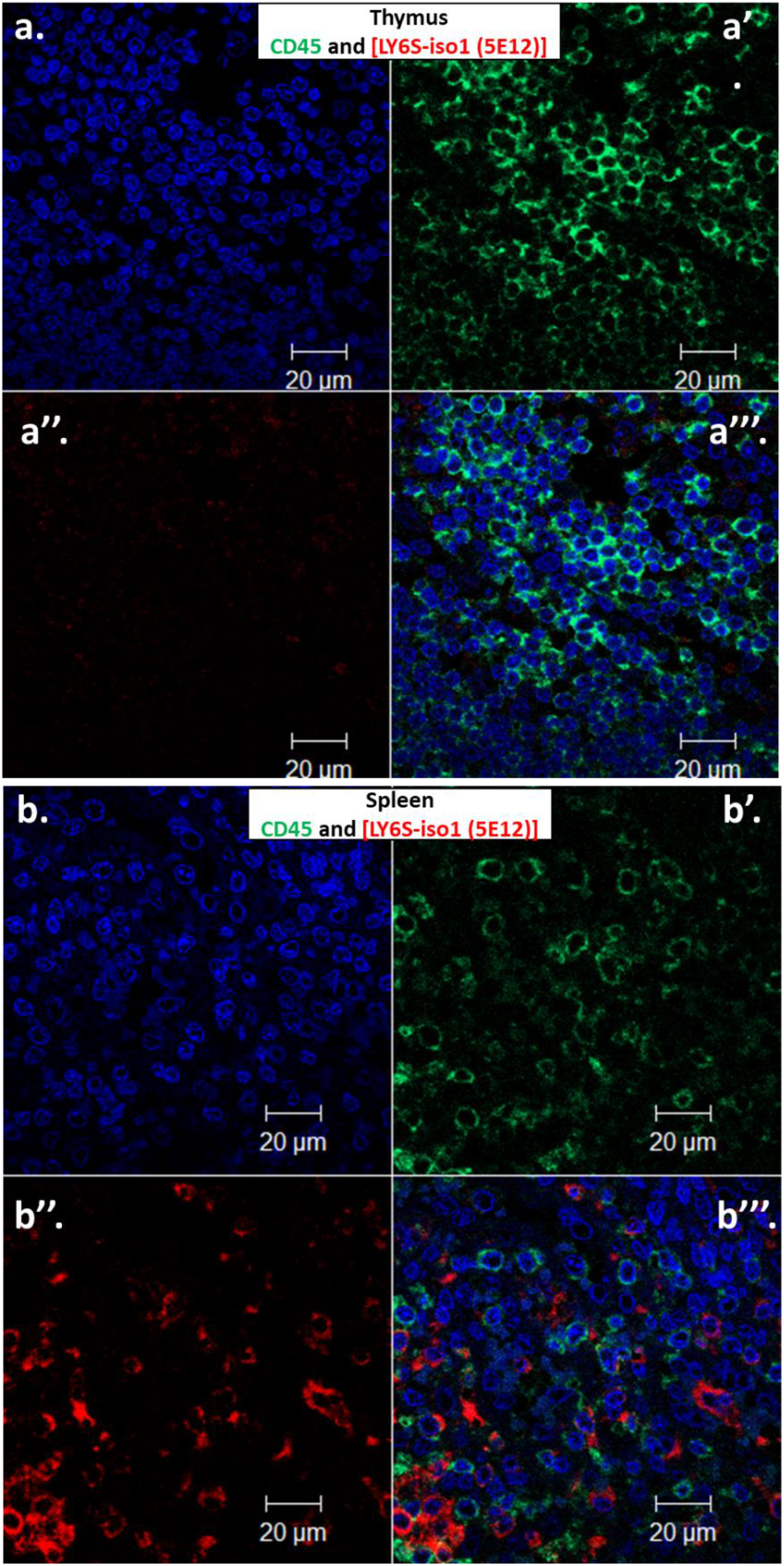
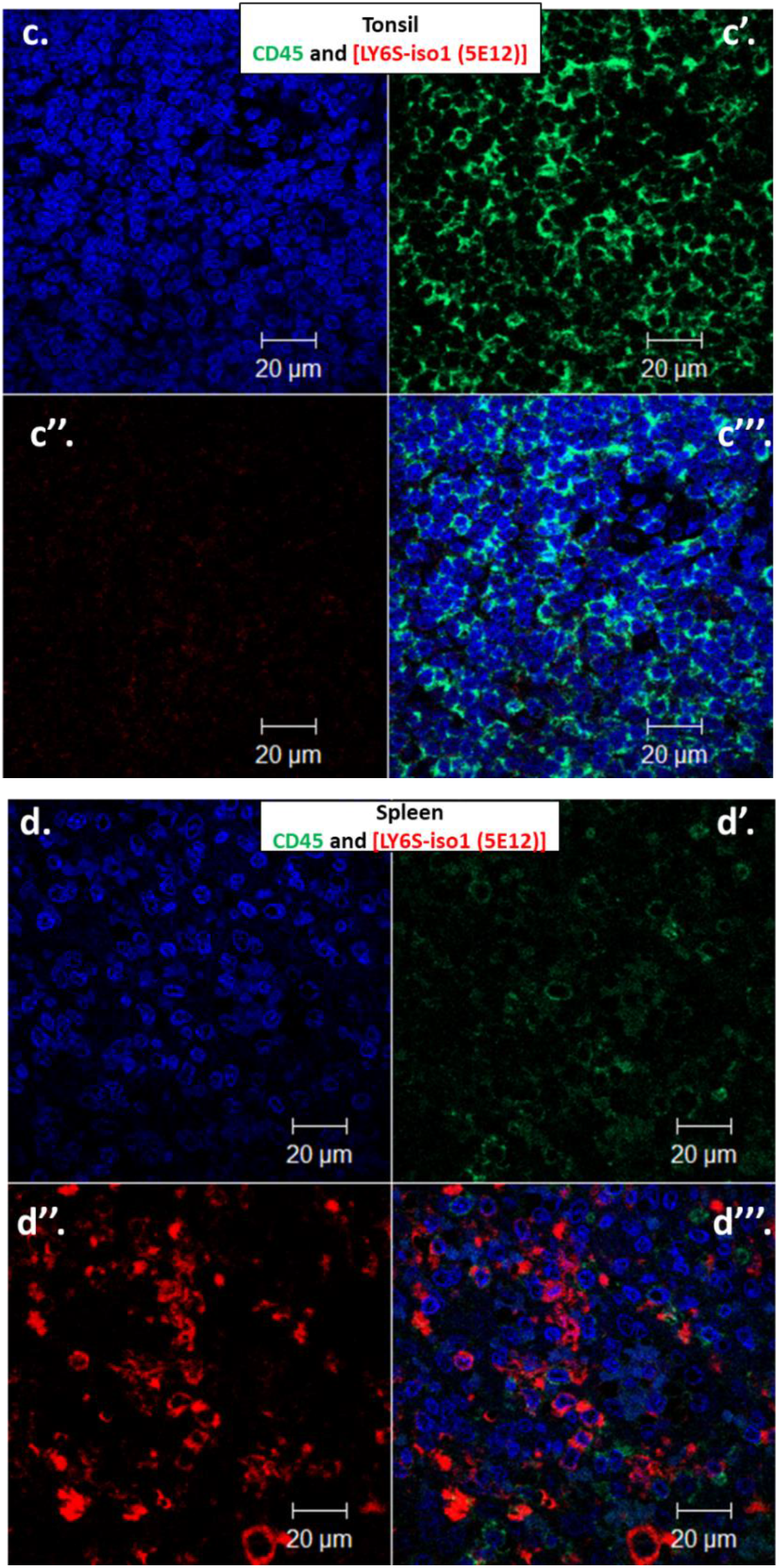
Co-immunofluorescent staining of LY6S-iso1 and CD45 in thymus, tonsil and spleen. Sections of thymus (panels a), spleen (panels b and d) and tonsil (panel c) were co-immunofluorescently stained for LY6S-iso1 (red staining, panels a”, b”, c” and d”) and CD45 (green staining, panels a’, b’, c’ and d’) or with DAPI (panels a, b, c and d). Merging of the stains is shown in panels a’’’, b’’’, c”’ and d’’’.

**Fig. S10.**
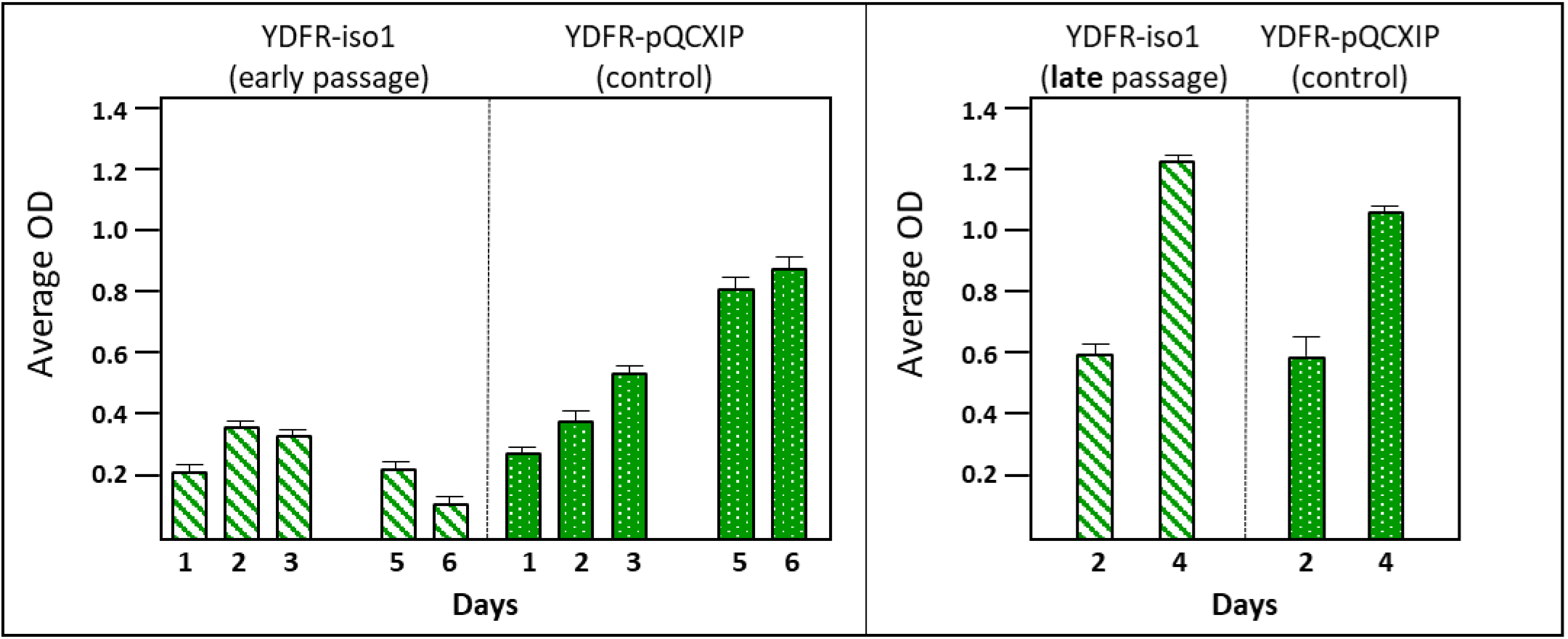
Growth of YDFR melanoma cells infected with LY6S-iso1. YDFR melanoma cells stably infected with control pQCXIP plasmid (YDFR-pQCXIP) or YDFR cells expressing LY6S-iso1 taken at an early passage or late passage [YDFR-iso1 (early passage) and YDFR-iso1 (late passage)] were seeded in wells of a 96-well culture plate (15,000 cells were seeded per well for the experiment at the left and 20,000 cells were seeded for the experiment at the right). Cell growth was monitored by the alkaline phosphatase assay. Note that in the 96 well format early passage YDFR-iso1 cells (at the left) initially grew very slowly, then remained stationary followed by loss of cell viability, whereas late passage YDFR-iso1 cells grew at a similar rate to the control YDFR cells (at the right).

**Fig. S11.**
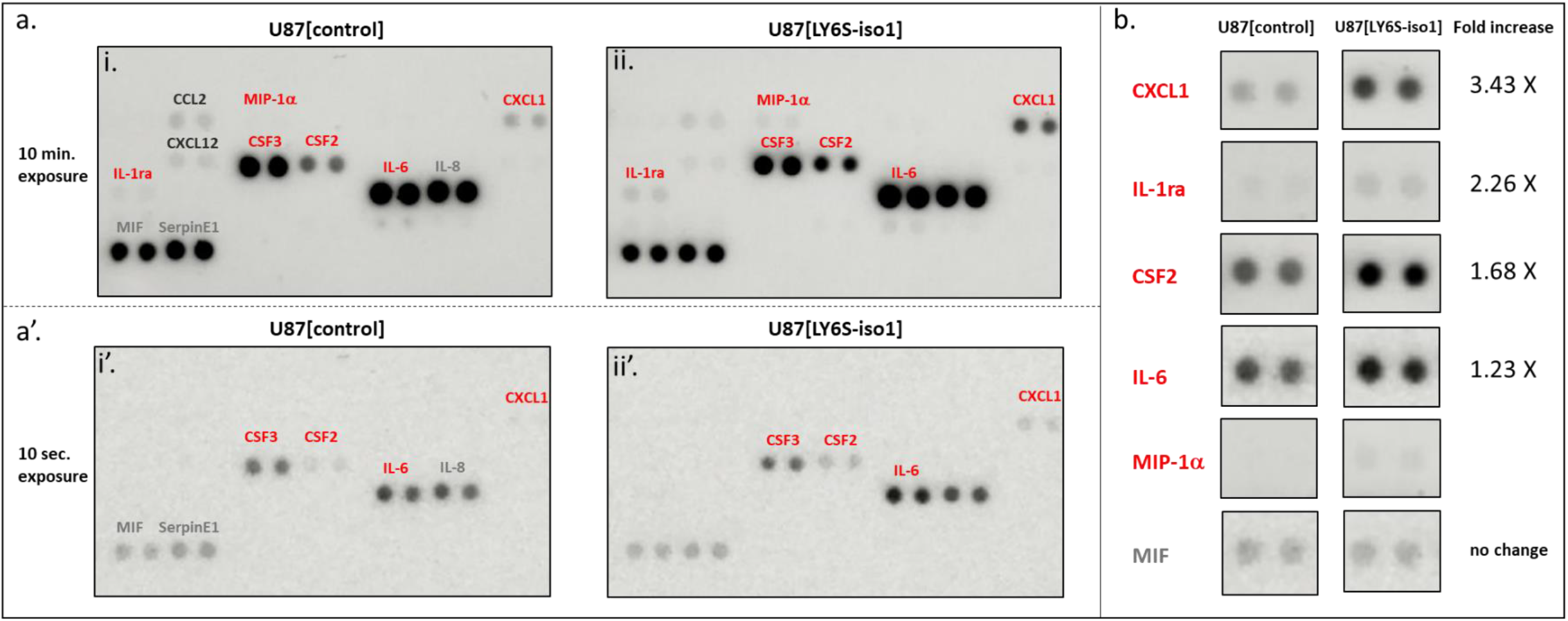
Profiling of selected cytokines/chemokines secreted by control and LY6S-iso1 expressing U87 glioblastoma cells. ***Panel a*-** Spent culture medium from U87 cells stably transfected with control pQCXIP plasmid (U87[control], panels a-i and a-i’) or from U87 cells stably transfected with plasmid coding for LY6S-iso1 (U87[LY6S-iso1], panels a-ii and a-ii’) were assayed for secreted cytokines and chemokines using a human cytokine/chemokine array. The probed arrays were exposed in a light imager either for 10 minutes (panels a-i and a-ii) or for 10 seconds (panels a-i’ and a-ii’). Cytokines/chemokines whose expression is increased in the LY6S-iso1 expressing cells are indicated in red fonts, whereas cytokines/chemokines whose expression remains unchanged are indicated by the grey fonts (note that CCL2 and CXCL12 appear to be slightly down regulated in the LY6S-iso1 expressing cells and are indicated with dark grey fonts). ***Panel b***-Side-by-side comparisons of the chemokines/cytokines upregulated in U87[LY6S-iso1] cells using optimal exposure times to demonstrate the upregulation. An internal control for equal loading of spent medium from both cell types was provided by MIF (lower two panels), that showed equivalent levels of expression in both control and [LY6S-iso1 expressing] cells. Quantitation for the fold increase in the secreted cytokines was performed by ImageJ analyses-as no signal was seen in the control cells for MIP-1alpha, its fold increase in LY6S-iso1-expressing cells could not be calculated.

**Fig. S12.**
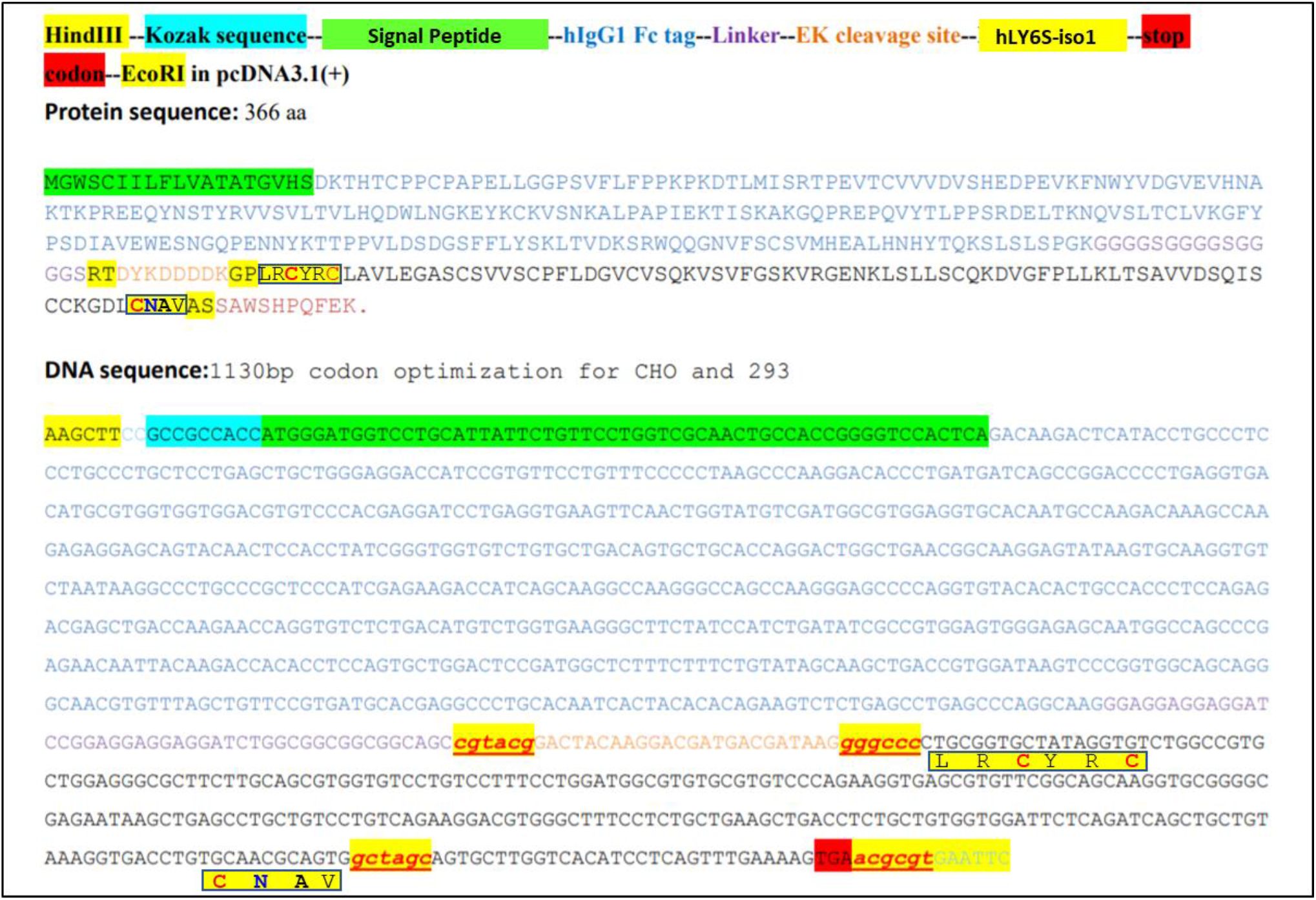
Expression of hFc-LY6S fusion protein. The make-up of the insert coding for the hFc-LY6S fusion protein, the amino acid sequence of the resultant fusion protein and the DNA sequence of the insert are shown in the top, second from top and bottom sections, respectively. This insert was cloned into the pcDNA3.1 plasmid, and stable HEK293 transfectants were generated that secrete into the cell culture medium the hFc-hLY6S-iso1 fusion protein. The amino acid sequences at the N- and C-termini of the LY6S-iso1 protein are indicated by the boxed sequences LRCYRC and CNAV, respectively.

**Table S1.**
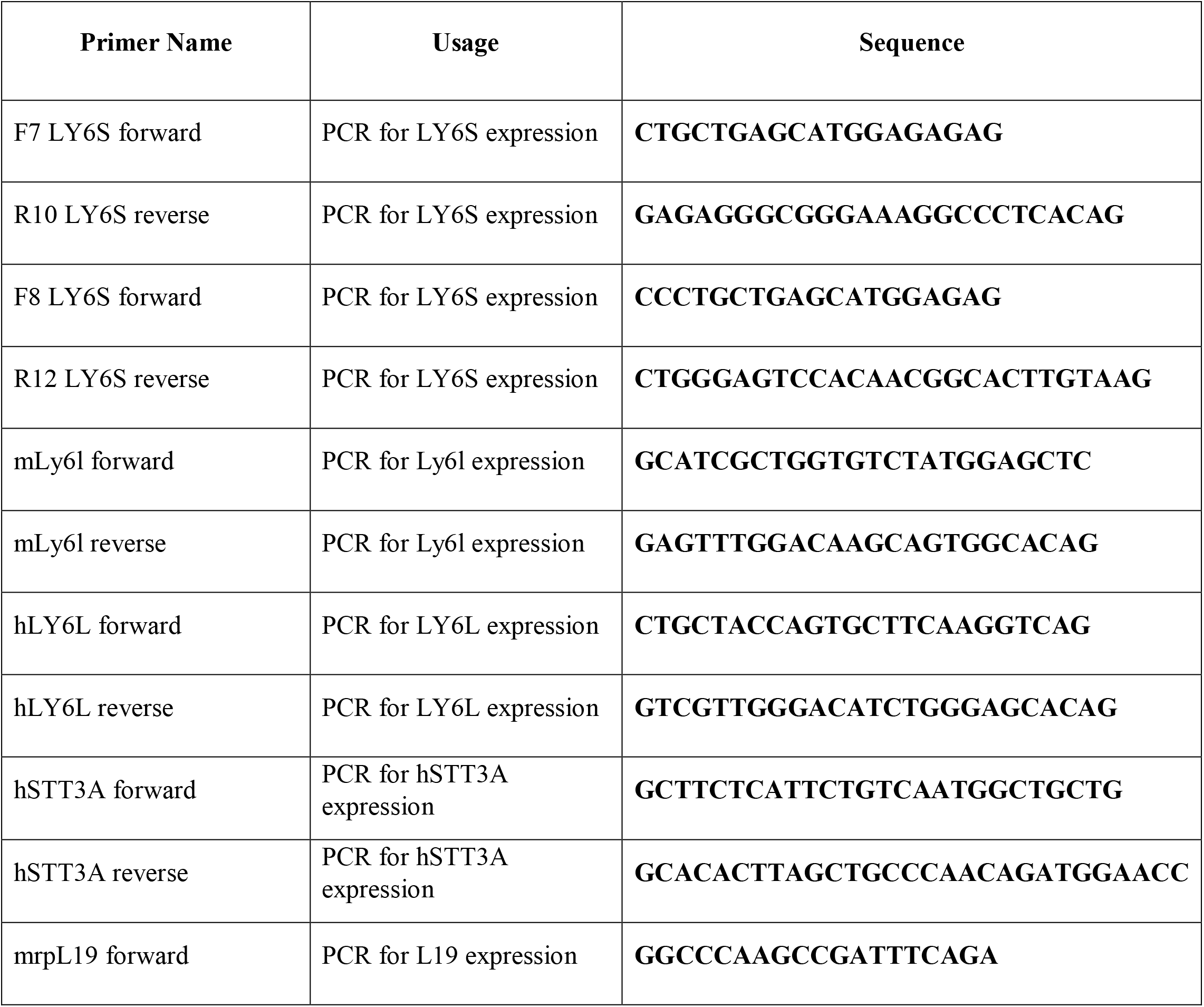
Oligonucleotides used in this study.

**Table S2.**
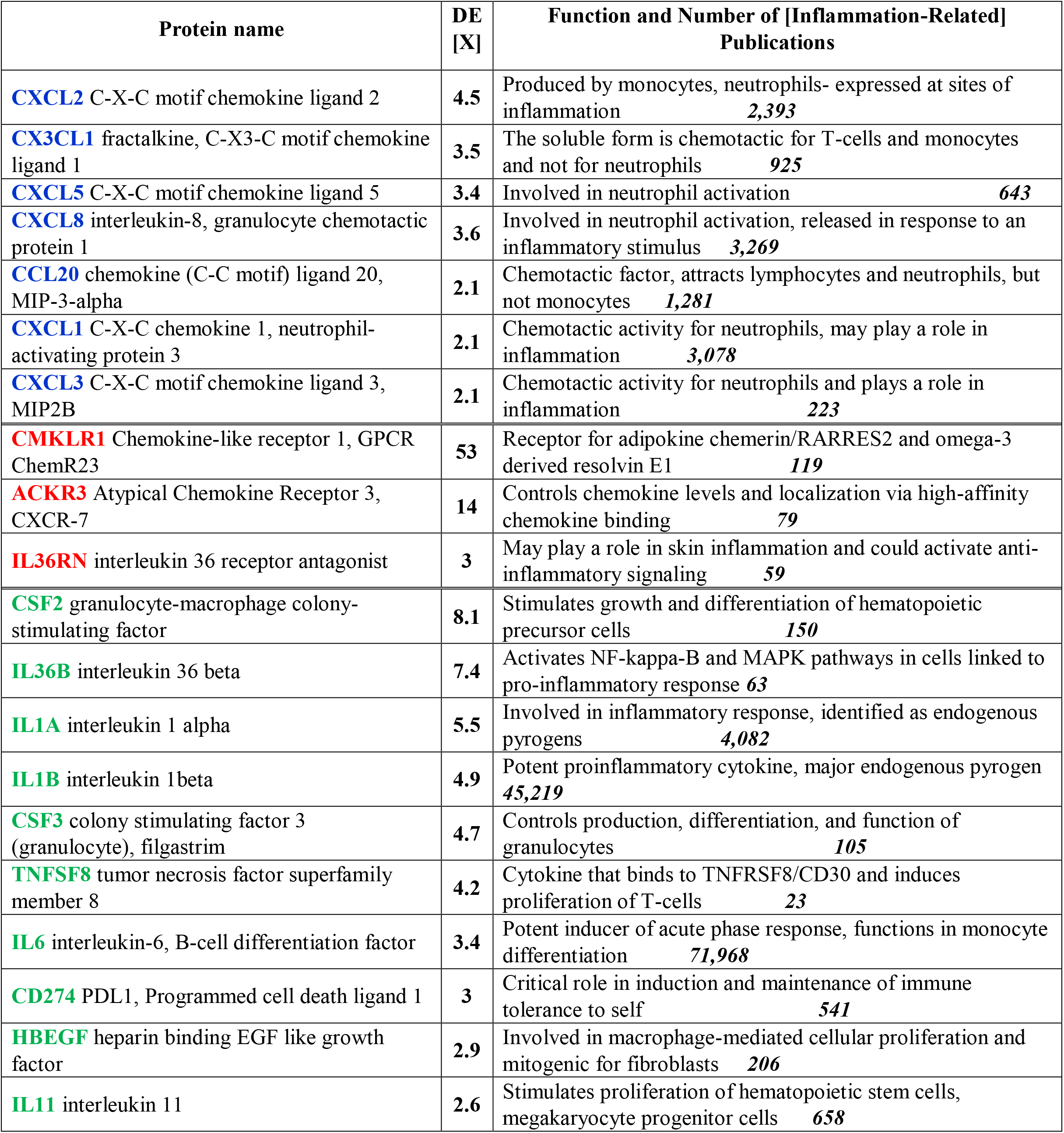
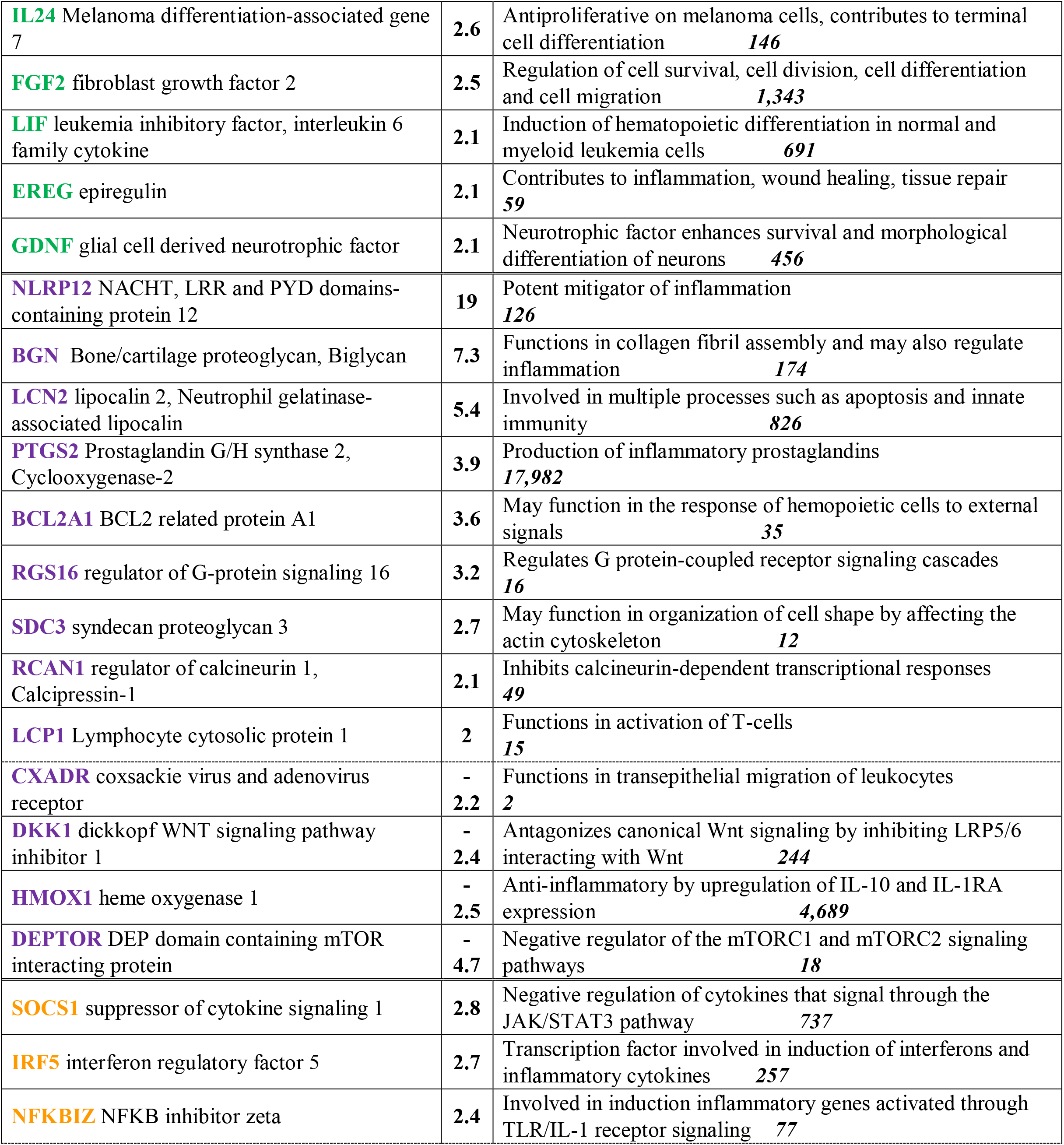
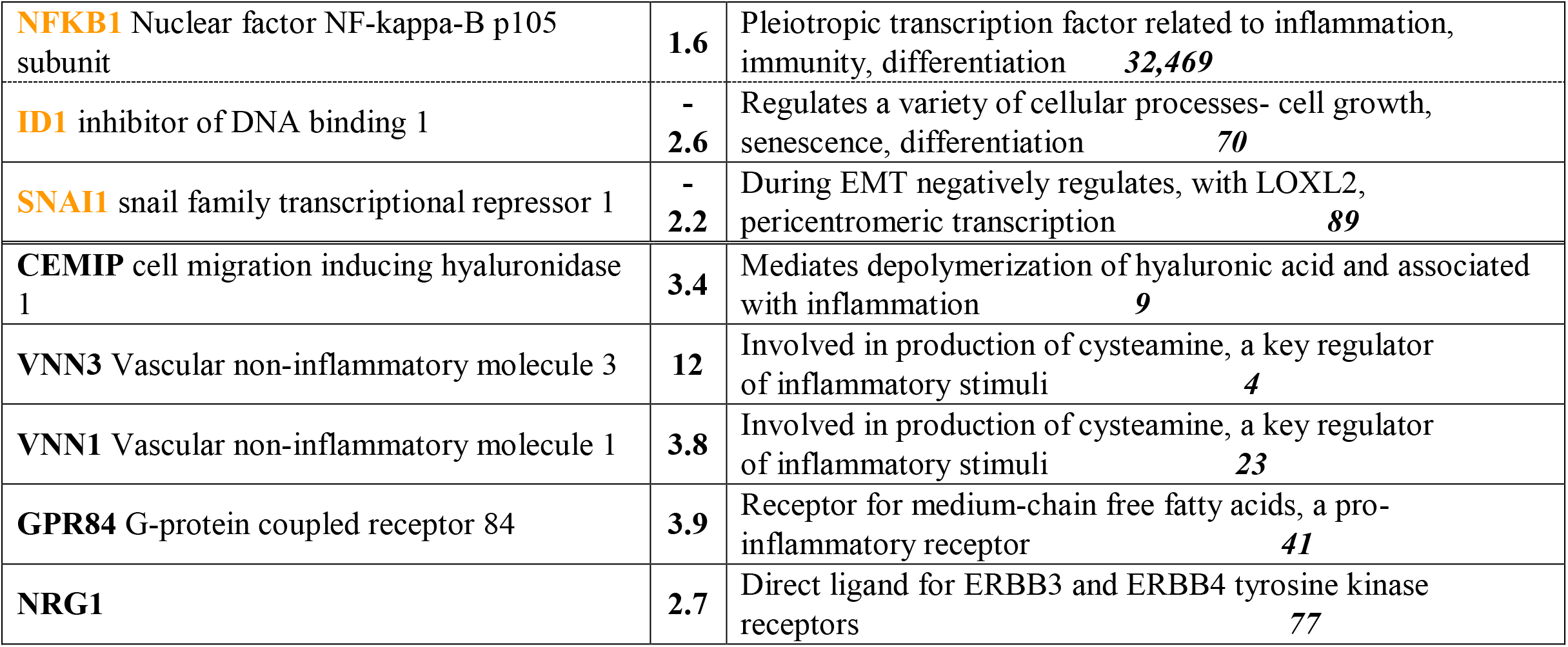
Names of proteins (‘Protein Name’) coded by genes differentially expressed (DE) in U87 human glioblastoma cells expressing LY6S-iso1 as compared to control U87 cells and the fold change in expression (‘DE X’). The column at right (‘Function and Number of Inflammation-Related Publications’) gives a brief synopsis of protein function and the number of publications relating to the protein (in bold italics at far right) retrieved with a PubMed search using as search terms [‘protein name’ and (inflammation OR inflammatory)].

**Table S3.**
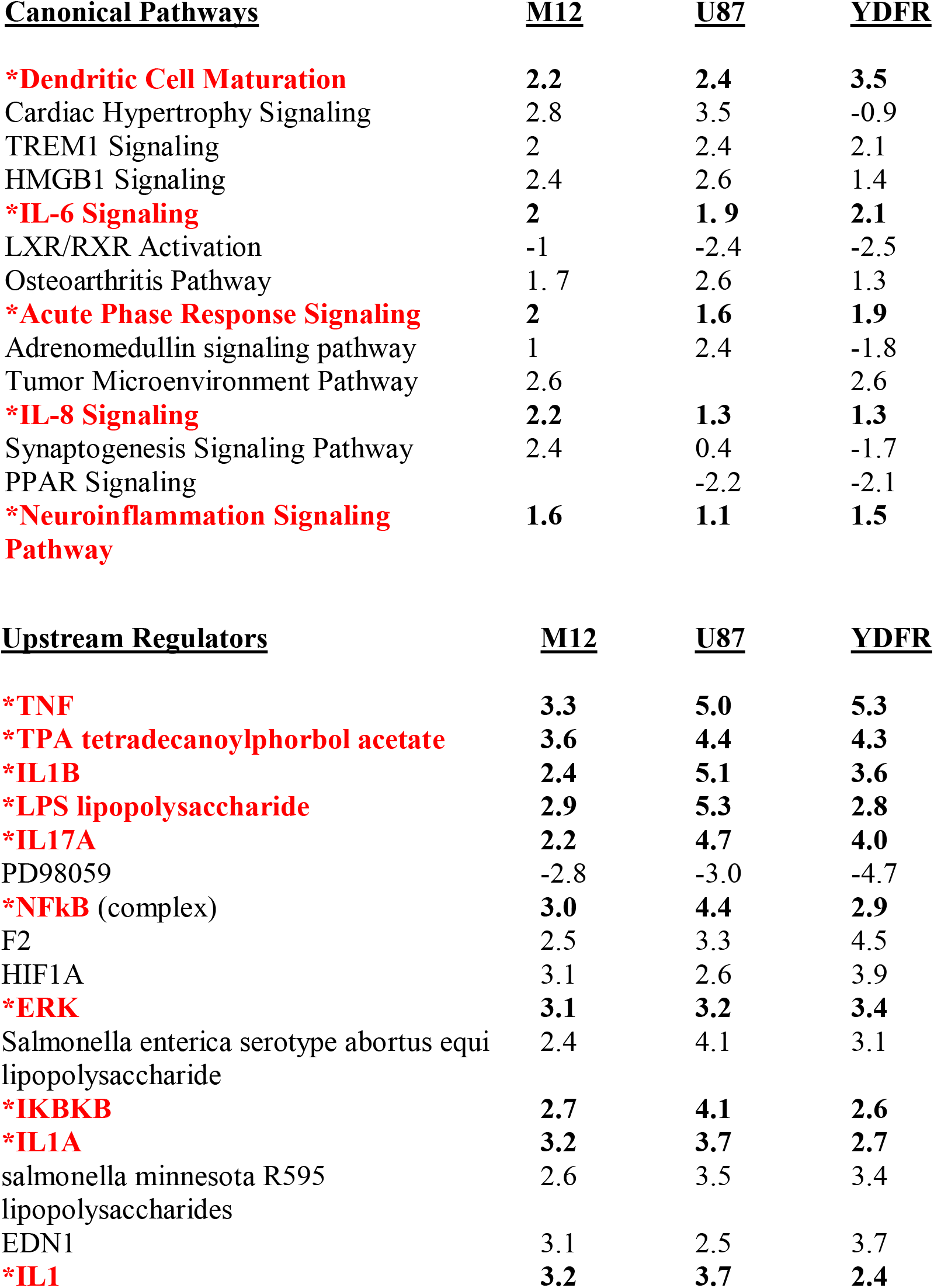
Canonical Pathways and Upstream Regulators. (upper and lower tables, with the z- scores shown) associated with expression of the LY6S-iso1 protein in M12, U87 and YDFR cells, as identified by the Ingenuity Pathway Analysis and by the IPA-derived Upstream Regulator Analysis.

